# Chemical Imaging Reveals Diverse Functions of Tricarboxylic Acid Metabolites in Root Growth and Development

**DOI:** 10.1101/2022.10.04.510836

**Authors:** Tao Zhang, Sarah E. Noll, Jesus T. Peng, Amman Klair, Abigail Tripka, Nathan Stutzman, Casey Cheng, Richard N. Zare, Alexandra J. Dickinson

**Author notes:** These authors contributed equally to this work (Tao Zhang, Sarah E. Noll). **Author contributions** A.J.D. and R.N.Z. conceived and supervised the project. T.Z. designed and performed the biological experiments. S.E.N. designed and performed the mass spectrometry experiments. J.P., A.K., A.T., N.S. and C.C. assisted with experimental work. T.Z., S.E.N., R.N.Z. and A.J.D. wrote the manuscript with the help of the other authors. **Competing Interest Statement:** The authors declare no competing interest.

## Abstract

Understanding how plants grow is critical for agriculture and fundamental for illuminating principles of multicellular development ^1^. Here, we apply chemical mapping of the developing maize root using desorption electrospray ionization mass spectrometry imaging (DESI-MSI) ^2^. This technique reveals a range of small molecule distribution patterns across the gradient of stem cell differentiation in the root. To understand the developmental logic of these patterns, we examined tricarboxylic acid (TCA) cycle metabolites. In both Arabidopsis and maize, TCA metabolites are enriched in developmentally opposing regions, suggesting that stem-cell specific TCA metabolite localization may be conserved in evolutionarily divergent species. We find that these metabolites, particularly succinate, aconitate, citrate, and α-ketoglutarate, control root development in diverse and distinct ways. Critically, the effects of metabolites on stem cell behavior can be independent of their canonical role in ATP production. These results present new insights into development and suggest practical means for controlling plant growth.

## Main

### DESI-MSI reveals small molecule distribution patterns along the developmental axis of the maize root

In the study of multicellular development, plant roots present a unique opportunity. The longitudinal axis of a root transitions through multiple stages of development, from the birth of new stem cells in the rapidly dividing meristem to formation of mature cells types in the differentiation zone. This permits the observation of continuous developmental gradients within a single slice of tissue (Fig. 1a) ^3^. Elucidation of mechanisms that regulate root growth have relied extensively on visualization of biomolecules along this developmental gradient. However, although small molecules are essential regulators of root development, it has remained a challenge to visualize them in their native context ^4^. Here, we apply desorption electrospray ionization mass spectrometry imaging (DESI-MSI) ^2^ to map the identities and relative intensities of molecules along the developmental gradient of the root.

**Fig. 1.**
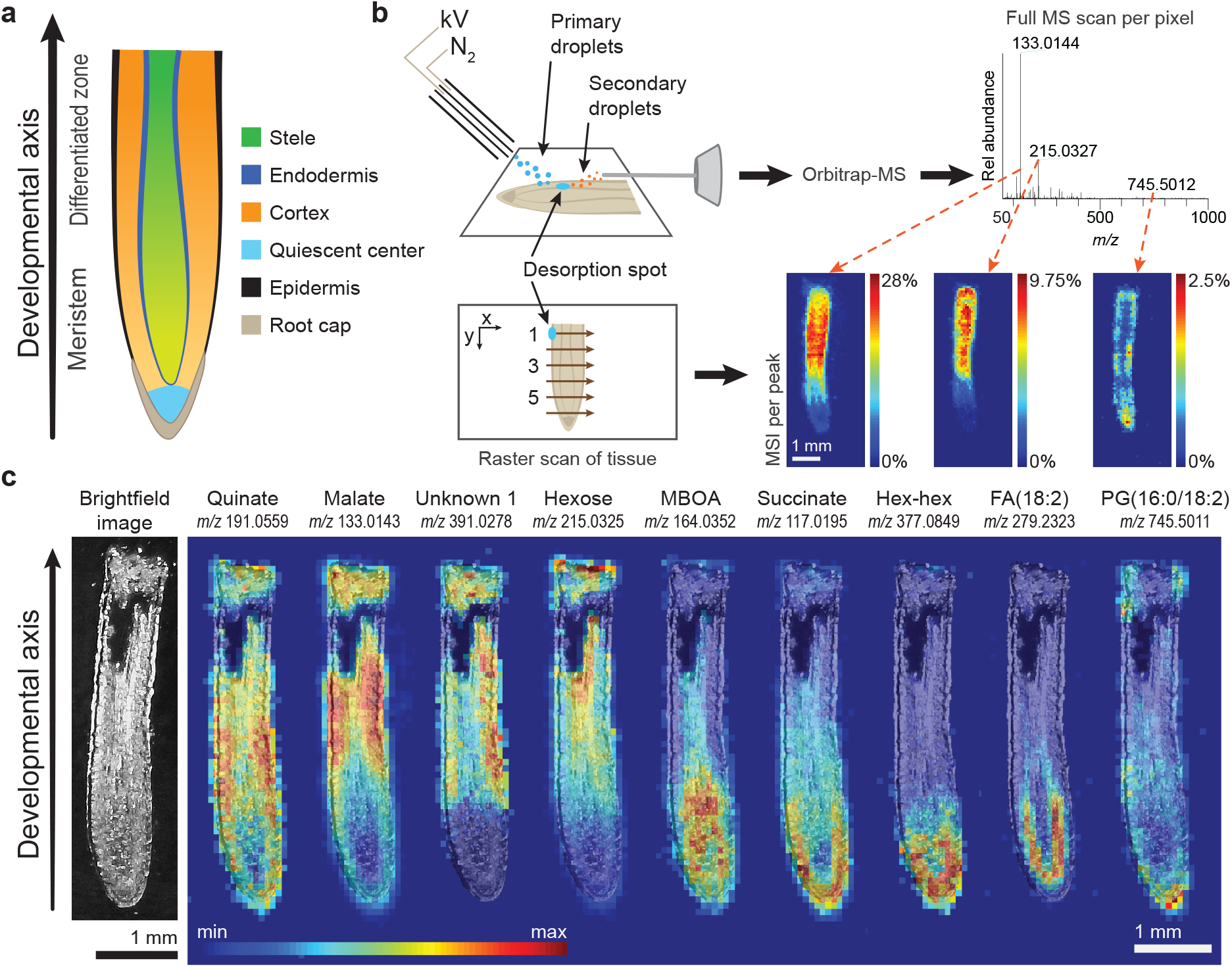
DESI-MSI captures the distinct distribution patterns of endogenous molecules within the developing maize root. **a**, Maize root tip with developmental zones and tissue types. **b**, Schematic of DESI-MSI shows the mechanism of ambient desorption and ionization by continuous spray of charged droplets, where one full scan is captured per pixel. The desorption spot is moved in a raster scan across the tissue to produce a pixelated map. Each peak in the full spectrum corresponds to one MS image. **c**, Brightfield image of a maize root section prior to DESI imaging. DESI-MS images show that certain metabolites and lipids are most intense in distinct tissues and developmental zones. MS images are overlaid with the brightfield image. The maximum intensity of each ion as a percentage of total ion current is: quinate, 1.40%; malate, 2.18%; unknown 1, 1.13%; hexose, 12.3%; 6-methoxy-2-benzoxazolinone (MBOA), 2.74%; succinate, 5.07%; hexose-hexose, 1.47%; fatty acid (18:2), 3.83%; and phosphatidylglycerol (16:0/18:2), 2.61%.

We imaged thin cryosections of maize (*Zea mays*) root tissue after careful adaptation of the DESI imaging technique to resolve the fine features of each root. DESI-MSI is an ambient ionization method that requires no sample preparation or derivatization. Briefly, a stream of charged droplets of a selected solvent mixture was directed at the surface of a root section, forming a thin film known as the desorption spot (Fig. 1b) ^2^. We modified the DESI sprayer and imaging parameters, which doubled the resolution of our method compared to traditional DESI-MSI (see Methods and Extended Data Fig. 1, 2, and 3). The droplet spray desorbs root material and carries it to the mass spectrometer for subsequent detection. By moving the desorption spot in a raster scan across the root section, a pixelated image is formed, where each pixel (picture element) contains a full mass spectrum. We coupled this technique to a high-resolution accurate mass (HRAM) orbitrap mass spectrometer for spatiochemical characterization.

MS images revealed a wealth of lipids, carbohydrates, and primary and secondary metabolites, including benzoxazinoids, a class of plant defense molecules (Fig. 1c, Supplementary Information Figs. 1-7) ^5–7^. Between 20 to 30 small molecules were localized per each developmental zone (Supplementary Figs. 8-10). We hypothesized that these enrichment patterns might predict developmental functions. Particularly striking were the observed differential gradients in the normalized intensity of TCA cycle metabolites along the developmental axis (Fig. 2a-c, Supplementary Figs. 11-15). In particular, succinate was most intense in the meristem. In contrast, aconitate, malate, and fumarate were typically enriched toward the region of root differentiation. We also observed an *m/z* signature consistent with both citrate and isocitrate, which are isomers and cannot be distinguished by mass spectrometry alone (Fig. 2a, Extended Data Table 1). To understand the developmental reasoning behind differential small molecule distribution patterns, we focused subsequent studies on determining whether individual TCA metabolites have specific effects on stem cell behavior.

**Fig. 2.**
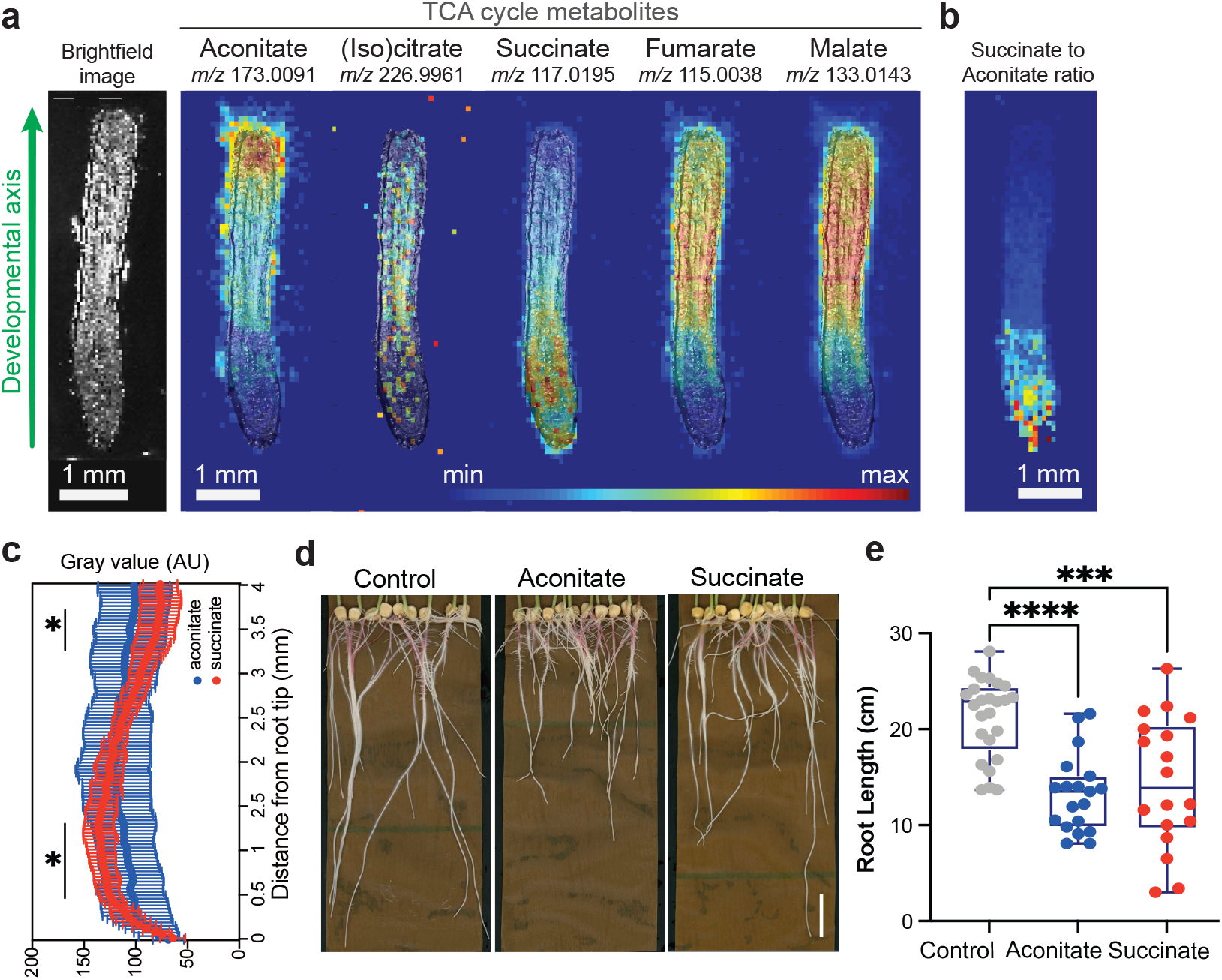
The TCA cycle metabolites are differentially distributed along the developmental axis of the root. **a**, DESI-MSI images of the detectable TCA cycle metabolites overlaid with a brightfield image captured prior to MS imaging. The maximum ion intensity as % TIC is different for each ion and is: aconitate, 2.99%; citrate/isocitrate, 0.271%; succinate, 3.12%; fumarate, 4.55%; and malate, 28.0%. **b**, MS image of the intensity ratio of succinate to aconitate. Maximum ratio is 10.9 (deep red). **c**, Normalized intensity of aconitate and succinate along the root axis. The data are presented as mean ± s.d. (n=10). Asterisks indicate statistical significance (*p < 0.05). **d-e**, Maize root phenotypes under 10 mM aconitate and succinate treatment, respectively. Scale bars, 3 cm. For the boxplots, the central line indicates the median, the bounds of the box show the 25th and 75th percentiles and the whiskers indicate 1.5× interquartile range. Asterisks indicate statistical significance by one-way ANOVA (***p < 0.001, ****p < 0.0001).

### TCA metabolites cause diverse developmental phenotypes in maize and Arabidopsis

We applied exogenous TCA metabolites to maize to explore the roles of these compounds in maize root development and growth. We found 10 mM aconitate, succinate, and malate inhibited primary root length (Fig. 2d-e, Extended Data Fig. 4) compared to control plants. Broadly, except for aconitate treatment, which increased citrate/isocitrate levels, none of these treatments led to significant changes in the overall profile of TCA metabolites compared to control plants (Extended Data Fig. 4e), suggesting that the pathway is not highly sensitive to influx of these metabolites.

To delve more deeply into the developmental functions of TCA metabolites in the root, we characterized their effects on the tractable model plant, Arabidopsis. Arabidopsis roots, typically ∼100 μm in width, are too small to image with our current DESI-MSI capabilities; so to explore whether TCA metabolites have developmental distribution patterns in Arabidopsis, we used previously published RNA-Seq data sets to calculate the expression levels of TCA biosynthesis genes in the meristem and maturation zones (Fig. 3a). We found spatially distinctive distribution patterns of several classes of TCA biosynthesis genes (Fig. 3a), which match localization signatures identified in a separate study of Arabidopsis roots ^8^. In particular, *ACONITASE* (*ACO*) and *SUCCINATE-COENZYME A LIGASE* (*SCL*) gene expression levels in Arabidopsis correlated with the metabolite distributions observed in maize roots, suggesting that the developmental distribution of TCA metabolites is conserved across evolutionarily divergent plants.

**Fig. 3.**
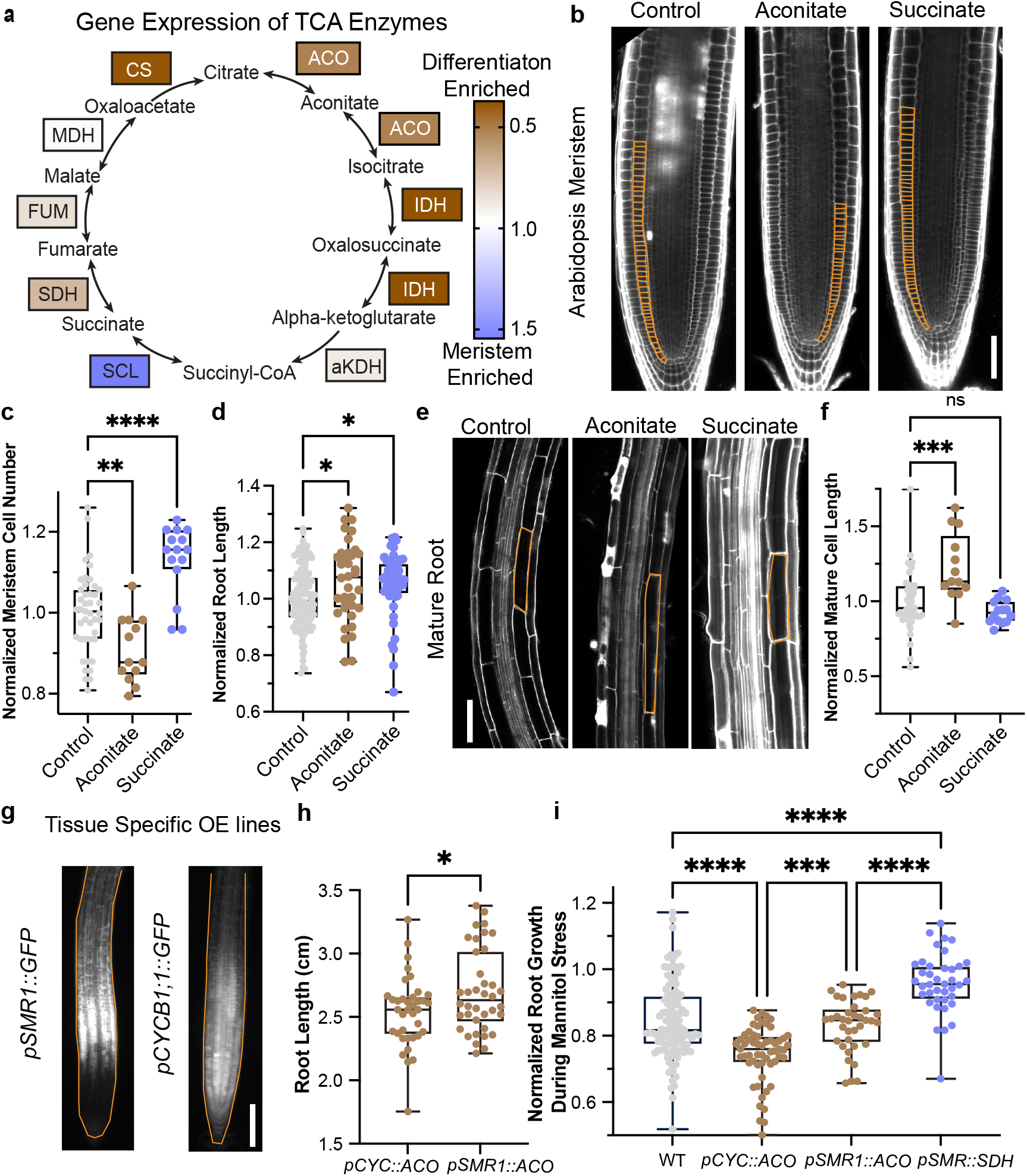
Succinate and aconitate have opposing effects on meristem cell divisions in Arabidopsis. **a**, Enrichment of the expression levels (log[3xFPKM]) of TCA biosynthesis genes in the root meristem. Data were quantified from Li S. et al. ^9^. Enrichment was calculated by dividing the average expression in the meristem by the average expression in the differentiation zone. **b**, Confocal images of root meristems treated with 1 mM succinate or aconitate, respectively. Scale bar, 100 μm. **c-d**, Quantification of the effects of 1 mM treatments on the number of meristematic cells and total primary root length, normalized to the control. **e**, Images of representative mature root cells. Scale bar, 100 μm. **f**, Quantification of average cell length in the mature root, normalized to the control. **g**, Transcriptional reporters for SMR, a promoter specific to the early differentiation zone, and *CYCB1;1*, a meristem promoter. Scale bar, 150 μm. Root outlines are highlighted in orange. **h**,**i**, Quantification of primary root length in genetically engineered lines (CYC = *CYCB1;1*, ACO = *ACONITASE 1*, SDH = *SUCCINATE DEHYDROGENASE 1*) with and without mannitol stress. Data was normalized to the control treatment for each genotype. For the boxplots, the central line indicates the median, the bounds of the box show the 25th and 75th percentiles and the whiskers indicate 1.5× interquartile range. Asterisks indicate statistical significance by one-way ANOVA (*p < 0.05, **p < 0.01, ***p < 0.001, ****p < 0.0001).

Because most of the TCA biosynthesis genes are knockout-lethal, we chose to apply exogenous TCA metabolite treatments to determine if these compounds induce changes in development. We identified distinct and concentration-dependent effects of several TCA metabolites on root development (Fig. 3 and Extended Data Fig. 5). Treatment with each TCA metabolite generates different changes in root growth, formation of branching roots (lateral roots and anchor roots), and root hair development, which is a hallmark feature of tissue differentiation. For example, treatment with either α-ketoglutarate or citrate (buffered to a pH of 5.7) inhibits primary root growth (Extended Data Fig. 5). However, these treatments have opposite effects on root branching. Citrate decreases lateral root number, but increases anchor root growth, whereas α-ketoglutarate significantly increases lateral root number and has no effect on anchor roots. Overall, investigation of the full TCA metabolite pathway reveals numerous distinct functions in root development and growth. Due to the complexity of the phenotypes induced by TCA metabolites, we chose to focus on well-characterized root stem cell behaviors: 1) divisions in the root meristem, 2) cell elongation upon exiting the meristem, and 3) differentiation into root hairs.

### TCA metabolite treatments alter meristematic cell divisions

To determine how TCA metabolites affect root growth, we measured meristem activity using confocal microscopy of exogenously treated roots (Fig. 3b-c, Extended Data Fig. 5). Citrate, α-ketoglutarate, and aconitate reduce meristem cell numbers. Isocitrate and succinate have the opposite effect, increasing meristem cell numbers significantly. Aconitate was the only metabolite that increases cell elongation, resulting in longer differentiated cells (Fig. 3e-f). Together, these results demonstrate that TCA metabolites lead to distinct root phenotypes in stem cell divisions and growth. Furthermore, these phenotypes correlate with their localization in the root. For instance, meristem-localized succinate is a meristem growth promoter whereas aconitate, which is enriched in the differentiation zone, inhibits meristem growth and promotes cell elongation. These correlations indicate that meristem enrichment can be predictive of molecular functions in stem cell behavior.

To explore the roles of TCA metabolite distribution patterns with more precision, we investigated the effect of changing them genetically. We ectopically overexpressed aconitate and succinate biosynthesis genes in specific developmental zones. We chose the meristem promoter *pCYCB1;1* ^10^ and the differentiation zone promoter *pSMR1* ^11^ to drive *ACO1* or *SDH1*, respectively (Fig. 3g-i). We found that disrupting the normal expression patterns of TCA biosynthesis genes can cause significant changes in root growth. Overexpressing *ACO1* in the differentiation zone significantly increases root growth compared to when it is overexpressed in the meristem. This finding is consistent with exogenous treatment results that show that aconitate promotes elongation in differentiating cells and inhibits cell divisions in the meristem. To explore TCA distribution patterns in sensitized conditions, we treated the transgenic plants with 100 mM mannitol, an osmotic stress used to mimic water deprivation (Extended Data Fig. 6). We found that overexpressing *SDH1* in the differentiation zone promotes resistance to 100 mM mannitol stress compared to wild type (Fig. 3i). In contrast, overexpressing *ACO1* in the meristem decreases primary root length compared to wild type plants. These results indicate that succinate and aconitate distribution patterns are important for the root to respond normally to environmental changes and can be manipulated to improve stress resistance.

### ATP levels are not affected by TCA metabolite treatment

One question these experiments raised is whether the phenotypes we observe result from a change in cellular energy, as the TCA cycle is a major source of ATP production. To determine whether these treatments affect ATP balance in the roots, we analyzed the effects of TCA metabolites on a fluorescent ATP sensor ^12^. We found that the localization pattern and intensity of the ATP sensor was fairly stable in response to changing TCA cycle inputs (Extended Data Fig. 7). Significant changes were induced by oxaloacetate and α-ketoglutarate treatments, which both increased the intensity of the ATP sensor in the meristem. Isocitrate was the only metabolite tested that significantly increased the intensity of the ATP sensor in the meristem and differentiation zone. This is consistent with the positive effect that isocitrate has on cell divisions (Extended Data Fig. 5d). These results suggest that isocitrate may provide a useful new starting point for biotechnology focused on increasing root ATP production and growth. Interestingly, these results demonstrated that changes in ATP levels are not sufficient to fully understand the differential phenotypes caused by individual TCA metabolite treatments.

### TCA metabolites had differential effects on the balance of reactive oxygen species (ROS) in Arabidopsis roots

Previous work on Arabidopsis roots has demonstrated that the spatial balance between hydrogen peroxide (H_2_O_2_) and superoxide (O_2•−_) is important for regulating the transition between stem cell proliferation and differentiation in the root ^13–16^. The alterations to cellular redox states caused by increase in H_2_O_2_ and O_2•−_ levels have been linked to the inhibition and maintenance of meristem cell divisions, respectively. SDH is a mitochondrial source of ROS in plants and animals ^17,18^. To test whether TCA metabolite treatments, particularly succinate treatment, regulate growth through ROS, we measured H_2_O_2_ and O_2•−_ in the root (see Extended Data Text and Extended Data Fig. 8). We found that all eight of the TCA metabolites tested increase O_2•−_ in the lower meristem, upper meristem, and/or differentiation zone (Extended Data Fig. 8a-b). Succinate increases O_2•−_ in the lower meristem and differentiation zone, which is consistent with the role of O_2•−_ in promoting meristem divisions. However, given the general induction of O_2•−_ by the TCA metabolites, this result is not sufficient to explain the specific promotional effect of succinate in cell divisions. Succinate also significantly increases H_2_O_2_ levels in the lower meristem, which would be expected to inhibit growth. This result further indicates that succinate’s effect on ROS is not sufficient to fully understand its role in root growth. The only other TCA metabolite that alters the levels of H_2_O_2_ in the plant root at 1 mM concentration is α-ketoglutarate, which dramatically reduces H_2_O_2_ in the meristem. Notably, α-ketoglutarate treatment alters the balance of H_2_O_2_ and O_2•−_ in a way that would be predicted to stimulate cell divisions, but instead it causes the opposite effect. Overall, ROS levels were highly sensitive to the TCA metabolite treatments, but the changes we measured did not fully explain the different effects of treatments on the root meristem. A hypothesis consistent with these results and previous literature ^14,15^ is that the meristematic response to TCA metabolites is occurring through a different signaling pathway that overrides ROS regulation of meristem activity.

### TCA metabolites regulate root hair growth via H_2_O_2_

We observed consistent effects of the TCA metabolites on root hair development in maize and Arabidopsis. Root hairs are essential for collecting water, absorbing nutrients, promoting beneficial microorganismal interactions, and increasing carbon sequestration. Previous studies have shown that root hair development is tightly regulated by H_2_O_2 ^13^_. Specifically, increases in H_2_O_2_ levels result in longer, more dense root hairs, whereas decreasing H_2_O_2_ has the opposite effect.

In Arabidopsis, we found that 5 mM succinate treatment dramatically increases H_2_O_2_ levels throughout the root (Fig. 4a-b). In addition, succinate also increases root hair length and density compared to control seedlings (Fig. 4c, d-e), which is the expected phenotype based on its effect on H_2_O_2_. In roots treated with 5 mM α-ketoglutarate and 5 mM citrate, respectively, the difference between lateral root density is greatly enhanced compared to 1 mM treatment (Fig. 4c, f, Extended Data Fig. 5). However, both treatments significantly reduce H_2_O_2_ and lead to smooth roots with shorter and less dense root hairs (Fig. 4d-e). We also observed the same effect in maize under 10 mM TCA treatment (Extended Data Fig. 9). These results are also consistent with a model wherein TCA metabolites alter root hair development by changing H_2_O_2_ levels (Fig. 4g). Overall, these results suggest that optimizing TCA metabolite levels and distribution patterns is a potential new strategy for improving agricultural sustainability.

**Fig. 4.**
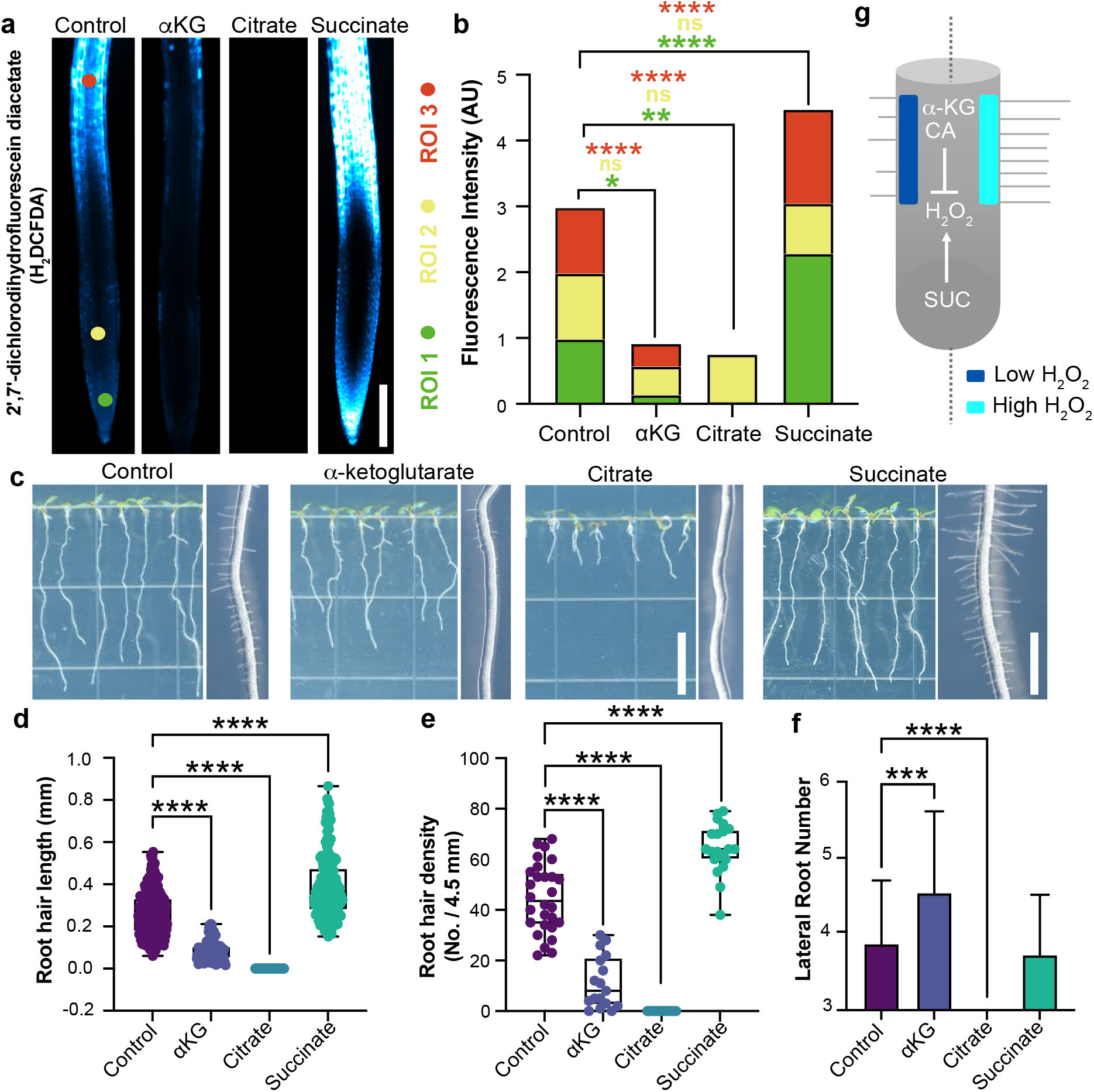
TCA metabolites alter ROS levels and root hair development. **a**, Images of H_2_DCFDA, a biological reporter for H_2_O_2_, in roots treated with 5 mM TCA metabolites. Scale bar, 200 μm. **b**, Quantification of the H_2_DCFDA fluorescence intensity in three regions of interest (ROIs). Data represent means ± s.d. (n = 15 plants). Asterisks indicate statistical significance by one-way ANOVA (*p < 0.05, **p < 0.01, ****p < 0.0001). **c**, Images of Arabidopsis roots treated with 5 mM TCA metabolites. Scale bar for primary root (left), 1 cm. Scale bar for root hair (right), 1 mm. **d-e**, Quantification of the effects of TCA metabolite treatments on root hair phenotypes. For the boxplots, the central line indicates the median, the bounds of the box show the 25th and 75th percentiles and the whiskers indicate 1.5× interquartile range. Asterisks indicate statistical significance by one-way ANOVA (***p < 0.001, ****p < 0.0001). **f**, Effect of TCA metabolites on lateral root number. Data represent means ± s.d. (n = 15 plants). Asterisks indicate statistical significance by one-way ANOVA (*p < 0.05, **p < 0.01, ***p < 0.001, ****p < 0.0001). **g**, A model for how succinate (SUC), citrate (CA), and α-ketoglutarate (αKG) influence root hair development through their effects on H_2_O_2_ levels.

## Discussion

TCA metabolites are increasingly being studied for their roles in signaling in a range of different organisms ^19–22^. However, analyzing TCA metabolite signaling has been highly challenging because most biosynthetic genes are essential and cannot be inhibited or knocked out without causing severe pleiotropic effects or lethality. In this study, spatial imaging of maize root chemistry using DESI-MSI revealed different patterns of TCA metabolite distributions along the axis of development. We found that exogenous treatments and tissue-specific genetic manipulation of TCA metabolites affected root growth in distinct ways. Certain phenotypes were readily predictable given the DESI-MSI localization data. For example, succinate and aconitate, which were enriched in the meristem and differentiation zone, respectively, had opposing effects on meristematic cell divisions. Tissue-specific overexpression of succinate and aconitate biosynthetic genes show significant effects on plant growth in control and stress conditions, which are consistent with their DESI-MSI localization patterns. However, other phenotypes were unexpected – including citrate’s promotion of anchor root growth, α-ketoglutarate’s promotion of lateral root density, and a range of effects from multiple metabolites on root hair development. We found that TCA-induced phenotypes were largely uncorrelated with changes in ATP localization or production. The exception was isocitrate treatment, which increased ATP levels and stimulated root growth. ROS levels, in contrast, are highly susceptible to TCA-cycle perturbations. The effects of TCA-cycle metabolites on ROS levels are consistent with the well-characterized regulatory roles of ROS in root hair development. Overall, this work suggests that multiple TCA metabolites have noncanonical roles in root development, providing interesting opportunities to improve our knowledge of how plant stem cells divide, differentiate, and perform organogenesis.

These results suggest a promising avenue for agricultural research, where a major challenge is applying the information that we learn in model species to genetically divergent crops. Control of TCA metabolites may be particularly amenable for engineering desirable phenotypes in diverse plants because the TCA cycle is so highly conserved. This approach may be especially important given current efforts to improve plant growth in the increasingly harsh environmental conditions caused by climate change.

## Methods

### Mass Spectrometry Materials

Dimethyl-formamide (certified ACS grade), acetonitrile (optima), methanol (HPLC grade) and water (HPLC grade) were purchased from Fisher Scientific (Waltham, MA). Fused silica capillaries were obtained from Polymicro Technologies, a subsidiary of Molex (Phoenix, AZ). 1/16” stainless steel T-junction and reducing unions was purchased from Swagelok (Solon, OH), while PEEK union assemblies, NanoTight sleeves, PFA tubing, and 1/16” stainless steel tubing were acquired from IDEX Health & Sciences (Oak Harbor, WA). Graphite ferrules were sourced from Restek (Bellefonte, PA). A syringe with a 22 gauge, point type 3 (blunt) removable needle was purchased from Hamilton (Reno, NV). Fisherbrand™ Superfrost Plus™ slides, Fisherbrand™ 5 mL sterile, single-use syringes, and 0.2 μm PES Thermo Scientific™ Nalgene™ Syringe Filters were purchased from Thermo Fisher Scientific (Waltham, MA).

### Plant growth and condition

For the Arabidopsis treatments, compounds were adjusted the pH to 5.7 and added to sterile 1/2X Murashige and Skoog (MS) plant growth media and adjusted the pH to 5.7 using potassium hydroxide. Seeds were sterilized using 20% bleach and sowed on the treated plant growth media. Plants were incubated in a 16h/8h day/night cycle growth chamber at 22 °C for 7-14 days prior to phenotyping. For the maize treatments, compounds were added to water and the pH was adjusted to 5.7 using potassium hydroxide. Maize seeds were sterilized using 20% bleach and grown in large size seed germination pouches (https://mega-international.com) with the treatment. Plants were incubated in a 16h/8h day/night cycle growth chamber at 28 °C night/21 °C day for 5 days prior to phenotyping.

### RNA seq analysis

RNA-seq data was analyzed using data published in “High resolution expression map of the Arabidopsis root reveals alternative splicing and lincRNA regulation.” ^9^. In this work, RNA was extracted and sequenced from three regions of the developing Arabidopsis root: the meristem, elongation zone, and differentiation zone.

### Constructs

The backbone of ectopic expression constructs is the plasmid *pHDE*, which was described previously ^23^. The promoter sequences of *pSMR1* and *pCYCB1* which are 1962bp and 1179bp respectively were obtained from genome DNA by PCR. The CDS sequence of *AtFUM1, ATCAO1, AtSDH1*, and *AtSSADH1* were obtained from Cdna which was transcribed from RNA. Those fragments with eGFP were assembled by Gibson assembly at the *Pme*I site. The plasmids were transformed into Arabidopsis Columbia plants by Agrobacterium-mediated floral dipping. Transgenic seedlings were selected on 1/2 X MS medium containing 16.7mg/L hygromycin.

### ROS staining

H_2_O_2_ was detected in roots using the H2DCFDA staining method. The roots from 7-day-old seedlings were incubated in H_2_O containing 50 µM H_2_DCFDA for 30 min under the dark condition. Then, the roots were rinsed with sterilized water three times. The fluorescent signals were observed with an Echo Revolution microscope (Echo). The excitation and emission wavelengths used for detection of the signals were 488 nm and 520 nm, respectively.

To detect superoxide, seedlings were stained for 2 min in a solution of 2.5 mM NBT in 50 mM phosphate buffer (pH 6.1) in the dark and then rinsed three times with distilled water ^24^. Images for NBT staining were obtained using a ×1 objective with a Nikon SMZ1270 Stereo microscope.

### Confocal images

For *A. thaliana*, most of the roots were stained by PI (Propidium iodide) and observed under a Zeiss LSM880 confocal microscope (PI, 561 nm; GFP, 488 nm), with thanks to Mark Estelle for allowing us to use it. Some of the roots were stained by PI and observed under a Leica SP8 confocal microscope (PI, 561 nm; GFP, 488 nm), thanks to the UCSD Microscopy Core --- NS047101. ATP sensor lines ^12^ were observed under a Leica SP8 confocal microscope (CFP, 439 nm; VENUS, 516 nm).

### HPLC-MS

For *A. thaliana*, each sample contained 10 whole roots which were collected 7 days after germination. For maize, each sample has a single root tip, 0.5 cm in length, which was collected 5 days after germination. We used sterilized deionized water as solvent. Root samples were flash frozen using liquid nitrogen and ground into a powder, which was then dissolved in water. The aqueous solution was placed in a 4 °C refrigerator overnight, after which it was centrifuged at 3000 rpm for 3 min. The supernatant was filtered using a syringe with a 0.22 μm membrane filter. The filtered solution was used for final analysis.

The parameters of the HPLC-MS method are as follows: Analysis was performed on a Thermo Scientific Vanquish UHPLC system coupled with an Orbitrap-MS (Orbitrap Elite, Thermo Scientific) with a heated electrospray ionization source. Chromatographic separation was carried out on a Kinetex® C18 column (150 × 2.1 mm, 1.7 mm) with a Kinetex® C18 guard column maintained at 35 °C. The mobile phases were A (H_2_O/formic acid (FA), 1000/1, v/v) and B (acetonitrile (ACN)/FA, 1000/1, v/v) with the gradient program: 0-6 min, 1% B; 6-7 min, 1% B to 90% B; 7-9 min, 90% B; 9-9.01 min, 90% B to 1% B; 9.01-12 min, 1% B. MS parameters included sheath gas flow rate of 50 arbitrary units, auxiliary gas flow rate of 20 arbitrary units, spray voltage of 3.5 kV, capillary temperature of 350 °C, S-lens RF level of 30, and resolution of 30,000. Measured SRM transitions (*m/*z 133 → 115, 173 → 85, 145 → 101, 191 → 111, 117 → 73, 131 → 85, and 115 → 71).

### Maize Tissue Samples for DESI-MSI

Maize B73 seeds were prepared for germination by soaking in distilled water for four hours at room temperature. Tip caps were then excised using a scalpel, and the seeds were sterilized with a 20% bleach solution. Seeds were incubated on damp germination paper inside a plastic petri dish.

Two days after germination, the primary root tip was cut, flash-frozen in liquid nitrogen, and mounted in optimal cutting temperature (O.C.T.) compound. 20 μm-thick tissue sections were cut with a cryostat and thaw-mounted onto a glass slide. Slides were stored in a -80°C freezer until imaging. Prior to imaging, slides were thawed in a vacuum desiccator.

### Optical Microscopy Images for DESI-MSI

Brightfield microscopy images of tissue sections were acquired prior to DESI-MS imaging at 250x magnification using a Pluggable USB 2.0 Digital Microscope with LED illumination (Redmond, WA).

### DESI-MS Imaging

First DESI images of maize roots were acquired as described previously ^25,26^. However, the probe and imaging parameters poorly resolved lateral features of the root (Extended Data Fig. 1c). To address this, modifications to the DESI probe (Extended Data Fig. 1a-b, d) were made, along with adjustments to the pixel size and ion injection time. All images for this study were acquired with the new probe and parameters. The modified DESI-MS probe was lab-built and constructed from a 1/16” stainless-steel Swagelok T-junction. The nebulizer tip consisted of two concentric fused silica capillaries. The outer sheath gas capillary was 350/250 μm (O.D./I.D.), while the inner solvent capillary was 150/20 μm. The sheath gas capillary was stabilized with a NanoTight sleeve and stainless-steel ferrule, whereas the solvent capillary was secured with a graphite ferrule.

A histologically compatible solvent mixture of 1:1 (v/v) dimethyl-formamide:acetonitrile at a flow rate of 1 μL/min was used ^27^. This solvent mixture provided good signal to background ratios while also preserving tissue integrity. Various methanol:water combinations (ranging from 70-100% methanol by volume) were explored, but cryosections were much more sensitive to ablation by increased water content (data not shown). Nitrogen sheath gas at 140 psi was used to aid nebulization. Images were acquired in the negative ion mode with a spray voltage of - 5 kV applied at the syringe needle. The probe tip was angled at 56° from the horizontal plane, and positioned approximately 2 mm above the slide surface, and ∼6 mm away from the extended ion transfer tube.

Images were acquired using a custom-build moving stage coupled to a high-resolution LTQ Orbitrap XL mass spectrometer (Thermo Fisher). Resolving power was 60,000 with the orbitrap as the mass analyzer, scanning over the mass range 50-1000. The ion injection time was 500 ms per microscan, and the automatic gain control was off. One full scan corresponded to one pixel. The pixel size was 72.6 × 80 μm. The pixel height was determined by the vertical step size, and the pixel width was calculated from the cycle time per scan and the lateral speed of the moving stage. The capillary voltage was -65 V; the tube lens voltage was -120 V; and the capillary temperature was 275°C.

In order to reduce batch effects, the ten B73 root tissues were imaged over two sequential days. The DESI probe was set-up and the spray optimized for the first day of imaging. The sprayer remained in place through the second day of imaging for continuity. The orbitrap was calibrated the day before imaging to ensure mass accuracy.

### DESI-MSI Data Analysis

Spectra were acquired in Xcalibur 2.5.5 (Thermo Fisher Scientific), where each row of the image was a separate .raw file. The .raw files were converted and compiled into one .imzML file, as described previously ^25^. DESI-MS images were visualized in MSiReader 1.01 ^28,29^.All images were normalized to the total ion current (see Supplementary Fig. 16), and the jet color scheme was used, where dark blue is the least intense and red is the most intense. The *m/z* tolerance of the images was set to ±5 ppm to match the expected mass accuracy of the orbitrap. Brightfield microscopy images were overlaid onto the MS images using MSiReader. The transparency was adjusted in MSiReader, and the black background of the optical image was removed in Paint3D (Microsoft) so as not to overly obscure the MS data.

To calculate the average intensity of aconitate and succinate along the developmental axis, grayscale DESI-MS images of each ion were uploaded to FIJI (open source package, ImageJ) ^30^. All images were normalized to the total ion current and scaled to the same maximum intensity. A slice was drawn through the center of each root, and the average intensity across a width of ten pixels was calculated for the length of the root. The average slices from the ten root sections were then averaged and reported (Fig. 2b).

The average spectra of the meristem, elongation, and differentiation zones were obtained by averaging across one row of one tissue in each respective developmental zone (Extended Data Fig. 3a-d).

### DESI-MSI Resolution Calculations

Lateral resolution was calculated using the 80-20% rule, which has been previously described for MS imaging ^31–33^. Briefly, this method defines resolution as the distance over which the signal rises from 20 to 80% of the maximum at an edge. A single ion chromatogram is extracted for these calculations. Here, phosphatidylglycerol (16:0/18:2) was chosen for the resolution calculations based on its signal strength and proximity to the tissue edge. It should be noted that any biological gradient in the selected ion may broaden the calculated resolution.

Single ion chromatograms were exported to Microsoft Excel, where a straight line was fit to the rising- and falling-edge using measured data points (Extended Data Fig. 2a). The point-slope formula was then used to calculate the *x*-coordinates of the 20% and 80% intensities. The difference in these coordinates represents the lateral spatial resolution. We calculated the rising- and falling-edge resolution and found there to be no significant difference in the means at a significance level of 0.05 using a two-way Student’s *t*-test with equal variance (Extended Data Fig. 2b). There was much intra-tissue variability in the measured resolution (Extended Data Fig. 2c-d), which suggested a lack of correlation in calculated values from each tissue section. To account for this, we included data from four rows of each of the ten tissue sections in calculating the average for each edge, resulting in an n-value of 40 for the rising- and falling-edge, respectively. This encompasses the total number of calculated resolution values, including multiple per tissue section. Statistical analysis, box plots, and half-violin plots were constructed in OriginPro 2021. For additional detail, please see^26^.

### Identification of Molecules by MS/MS

To definitively identify peaks of interest, we used collision induced dissociation (CID) of tissue extract. Primary root tips, 0.5 cm in length, were excised from five-day old B73 maize shoots five days after germination. After patting dry, two root tips were placed in each Eppendorf tube on dry ice. Roots were vacuum desiccated for 18 minutes, and then ground with a plastic pestle. 350 μL of methanol were added, and then tubes were shaken for 45 minutes (Analog Vortex Mixer, Ohaus), after which they were spun down at 3300 rpm. The supernatant was removed and filtered through a 0.2 μm PES filter into a new Eppendorf tube. The filter was rinsed with an additional 100 μL of methanol. Tissue extracts were prepared on the same day as analysis and kept on dry ice until CID was performed.

For electrospray ionization (ESI), a commercial Ion Max source with heated ESI spray tip was used with a high-resolution Orbitrap Elite mass spectrometer (Thermo Fisher). The solution flow rate was set to 5 μL/min. The heater temperature was set to 70°C, the sheath and auxiliary gas glow rates to 12 AU, the capillary temperature to 325°C, and the S-Lens RF Level to 60%. Data was acquired in the negative ion mode with a spray voltage of -3 kV. The orbitrap was used as mass analyzer with a resolving power of 120,000. Normalized collision energy ranged from 15-30%. These parameters were used for all collected CID, except the fragmentation patterns of PG(16:0/16:0) and PG(16:0/18:2), which were obtained on a different day and instrument, as described previously^26^.

Collected fragmentation patterns were compared to database values obtained from METLIN ^34,35^, METLIN Gen2, LipidMaps ^36^,and PubChem ^37^, as well as previously published literature data ^6,38^. A table of major fragment peaks is provided in the Extended Data Table 1. Full fragmentation spectra are provided in the Supplementary Information (Supplementary Information Figs. 17-35).

## Extended Data Figure Legends

**Extended Data Fig. 1** | **DESI-MSI probe design and imagine parameters for improved spatial resolutiona**, Generic schematic of the DESI probe shows the coaxial capillary construction at the tip, where the outer capillary is for the nitrogen sheath gas and the inner capillary runs continuously from the spray tip backward to the solvent syringe. **b**, Photograph of the DESI imaging set-up for maize root acquisition. Details of the coaxial probe tip, pixel size, ion injection time, and sample DESI-MS images of fumarate are shown for the initial maize imaging attempts **(c)** and the modified probe and parameters used for all image acquisition in this study **(d)**. MS images are of fumarate with brightfield microscopy image overlay. The outer borders and holes of the brightfield image are outlined in pink to demonstrate the improved resolution with the modified set-up.

**Extended Data Fig. 2** | **Lateral resolution was determined using the 80-20% rulea**, Demonstration of the 80-20% rule using the extracted ion chromatogram of PG (16:0/18:2) from one row of one maize tissue. The distance over which the intensity rises from 20% to 80% of the maximum for each edge is the lateral resolution. **b**, Lateral resolution for the rising- and falling-edges of the ten tissue sections. Dashed lines and labels show the mean values, while whiskers are one standard deviation. The box encompasses the 25th-75th percentile. Four rows were measured for each tissue, providing n = 40 for each edge. A two-way Student’s *t*-test showed no significant difference in the means at a significance level of 0.05. To justify the inclusion of four rows per tissue, we show the intra-tissue variability in the measured lateral resolution for the rising- **(c)** and falling-edge **(d)** of the ten measured tissues (n=4 per tissue). Dark gray circle and line represent the mean value and one standard deviation for each tissue section. Measured values are colored diamonds. The half-violin plots were constructed with bin widths of 20 μm.

**Extended Data Fig. 3** | **Representative mass spectra from distinct developmental zones of the roota**, Brightfield microscopy image of the root section from which a representative spectrum of the elongation zone (**b**, pink), meristem zone (**c**, blue), and root tip (**d**, green) are shown. The lines in **(a)** indicate the rows that were averaged to produce the spectra in **(b-d)**.

**Extended Data Fig. 4** | **Maize is insensitive to TCA metabolites treatment compared to Arabidopsisa***-***d**, Root phenotypes in maize seedlings treated with 1 mM and 5 mM aconitate, fumarate, malate, and succinate, respectively. **e-g**, Root phenotypes in maize seedlings treated with 10 mM fumarate, malate, and succinate. Scale bar, 3 cm. h, Analysis of TCA metabolites using HPLC-MS shows the effect of 10 mM metabolite treatments on the normalized level of five TCA metabolites. Data represent means ± s.d. (n =3 biological replicates). Asterisks indicate statistical significance by one-way ANOVA (**p < 0.01, ****p < 0.0001). Source data are provided as a Source Data file. Data are presented as boxplots with each dot representing the datapoint of one biological replicate. For the boxplots, the central line indicates the median, the bounds of the box show the 25th and 75th percentiles and the whiskers indicate 1.5× interquartile range.

**Extended Data Fig. 5** | **Response of Arabidopsis root to TCA metabolite treatmentsa**, Exogenous 1 mM TCA metabolite treatments showed a different affection on plant root development. Scale bar, 1 cm. **b**, Confocal images of meristem and differentiation zone cells, respectively, under 1 mM TCA metabolite treatments. Scale bar, 50 µm. **c**, Normalized primary root length of 1 mM TCA metabolite treatments. **d**, Number of cells in the root meristem zone (MZ) after 1 mM TCA metabolite treatments. **e**, Average cell length of cells in the differentiation zone (DZ) after 1 mM TCA metabolite treatments. Data are presented as boxplots with each dot representing the datapoint of one biological replicate. For the boxplots, the central line indicates the median, the bounds of the box show the 25th and 75th percentiles and the whiskers indicate 1.5× interquartile range. Asterisks indicate statistical significance by one-way ANOVA (*p < 0.05, ***p < 0.001, ****p < 0.0001). **f**, Average lateral roots number (LRN) of 1 mM TCA metabolite treatments. Data represent means ± s.d. (n = 30 plants). Asterisks indicate statistical significance by one-way ANOVA (**p < 0.01). **g**, Anchor roots number (ARN) of 1 mM TCA metabolite treatments. Data represent means ± s.d. (n = 30 plants).

**Extended Data Fig. 6** | **Ectopic expression of TCA metabolites by tissue-specific genetic engineeringa**, Schematic of constructs for ectopic expression of TCA cycle genes. **b**, Transgenic lines of *pSMR1-SDH1-GFP* show increased growth in mannitol stress compared to wild type. Scale bar, 1 cm. **c**, Transgenic lines of *pCYCB1;1-ACO1-GFP* are more sensitive to salt stress compared to wild type. Scale bar, 1 cm. **d**, Normalized root length of transgenic plants in standard media, media without phosphate, and media with 50 mM NaCl. For the boxplots, the central line indicates the median, the bounds of the box show the 25th and 75th percentiles and the whiskers indicate 1.5× interquartile range. Asterisks indicate statistical significance by one-way ANOVA (**p < 0.01, ***p < 0.001, ****p < 0.0001). Source data are provided as a Source Data file.

**Extended Data Fig. 7**. **Response of ATP in Arabidopsis root to 1 mM TCA metabolite treatmentsa**, Merged images of a confocal fluorescence (Venus and CFP) image of ATP sensor under 1 mM TCA metabolite treatments. Scale bar, 100 µm. **b-c**, Normalized Venus/CFP intensity of ATP sensor under 1 mM TCA metabolite treatments. Data represent means ± s.d. (n = 15 plants). Asterisks indicate statistical significance by one-way ANOVA (*p < 0.05, **p < 0.01, ***p < 0.001, ****p < 0.0001). Different colors indicate different TCA metabolite treatments.

**Extended Data Figs. 8** | **ROS staining of 1 mM TCA metabolite treatments of Arabidopsisa**, Images showing NBT staining; Scale bar, 2 mm. **b**, Normalized stain intensity of NBT stain in three regions of interest (ROIs). **c**, Images showing H_2_DCFDA staining. Scale bar, 100 µm. **d**, H_2_DCFDA with 1 mM TCA metabolite treatments in three regions of interest (ROIs). Data represent means ± s.d. (n = 15 plants). Asterisks indicate statistical significance by one-way ANOVA (*p < 0.05, **p < 0.01, ***p < 0.001, ****p < 0.0001).

**Extended Data Fig.9** | **Root hair phenotype of 10 mM TCA metabolite treatments in maizea**, Root hair phenotype of 10 mM α-ketoglutarate, citrate, and succinate treatment in maize. Scale bar, 0.15 mm. **b**, Normalized root hair density under different treatments. **c**, Root hair length under different treatments. For the boxplots, the central line indicates the median, the bounds of the box show the 25th and 75th percentiles and the whiskers indicate 1.5× interquartile range. Asterisks indicate statistical significance by one-way ANOVA (**p < 0.01, ****p < 0.0001).

## References

1. Schmidhuber, J. & Tubiello, F. N. Global food security under climate change. Proceedings of the National Academy of Sciences 104, 19703–19708 (2007).

2. Wiseman, J. M., Ifa, D. R., Song, Q. & Cooks, R. G. Tissue Imaging at Atmospheric Pressure Using Desorption Electrospray Ionization (DESI) Mass Spectrometry. Angewandte Chemie International Edition 45, 7188–7192 (2006).

3. Benfey, P. N. & Scheres, B. Root development. Curr Biol 10, R813–815 (2000).

4. Moussaieff, A. et al. High-resolution metabolic mapping of cell types in plant roots. Proceedings of the National Academy of Sciences 110, E1232–E1241 (2013).

5. Zhou, S., Richter, A. & Jander, G. Beyond Defense: Multiple Functions of Benzoxazinoids in Maize Metabolism. Plant Cell Physiol 59, 1528–1537 (2018).

6. de Bruijn, W. J. C., Vincken, J.-P., Duran, K. & Gruppen, H. Mass Spectrometric Characterization of Benzoxazinoid Glycosides from Rhizopus-Elicited Wheat (Triticum aestivum) Seedlings. J. Agric. Food Chem. 64, 6267–6276 (2016).

7. Hieta, J.-P., Sipari, N., Räikkönen, H., Keinänen, M. & Kostiainen, R. Mass Spectrometry Imaging of Arabidopsis thaliana Leaves at the Single-Cell Level by Infrared Laser Ablation Atmospheric Pressure Photoionization (LAAPPI). J. Am. Soc. Mass Spectrom. 32, 2895–2903 (2021).

8. Tang, M. et al. A genome-scale TF–DNA interaction network of transcriptional regulation of Arabidopsis primary and specialized metabolism. Mol Syst Biol 17, (2021).

9. Li, S., Yamada, M., Han, X., Ohler, U. & Benfey, P. N. High-Resolution Expression Map of the Arabidopsis Root Reveals Alternative Splicing and lincRNA Regulation. Developmental Cell 39, 508–522 (2016).

10. Colón-Carmona, A., You, R., Haimovitch-Gal, T. & Doerner, P. Spatio-temporal analysis of mitotic activity with a labile cyclin–GUS fusion protein. The Plant Journal 20, 503–508 (1999).

11. Yi, D. et al. The Arabidopsis SIAMESE-RELATED cyclin-dependent kinase inhibitors SMR5 and SMR7 regulate the DNA damage checkpoint in response to reactive oxygen species. Plant Cell 26, 296–309 (2014).

12. De Col, V. et al. ATP sensing in living plant cells reveals tissue gradients and stress dynamics of energy physiology. eLife 6, e26770 (2017).

13. Dunand, C., Crèvecoeur, M. & Penel, C. Distribution of superoxide and hydrogen peroxide in Arabidopsis root and their influence on root development: possible interaction with peroxidases. New Phytologist 174, 332–341 (2007).

14. Tsukagoshi, H., Busch, W. & Benfey, P. N. Transcriptional Regulation of ROS Controls Transition from Proliferation to Differentiation in the Root. Cell 143, 606–616 (2010).

15. Yamada, M., Han, X. & Benfey, P. N. RGF1 controls root meristem size through ROS signaling. Nature 577, 85–88 (2020).

16. Zhou, L. et al. Exogenous hydrogen peroxide inhibits primary root gravitropism by regulating auxin distribution during Arabidopsis seed germination. Plant Physiol Biochem 128, 126–133 (2018).

17. Jardim-Messeder, D. et al. Succinate dehydrogenase (mitochondrial complex II) is a source of reactive oxygen species in plants and regulates development and stress responses. New Phytologist 208, 776–789 (2015).

18. Quinlan, C. L. et al. Mitochondrial complex II can generate reactive oxygen species at high rates in both the forward and reverse reactions. J Biol Chem 287, 27255–27264 (2012).

19. Martínez-Reyes, I. & Chandel, N. S. Mitochondrial TCA cycle metabolites control physiology and disease. Nat Commun 11, 102 (2020).

20. Detraux, D. & Renard, P. Succinate as a New Actor in Pluripotency and Early Development? Metabolites 12, 651 (2022).

21. Ko, S. H. et al. Succinate promotes stem cell migration through the GPR91-dependent regulation of DRP1-mediated mitochondrial fission. Sci Rep 7, 12582 (2017).

22. Grimolizzi, F. & Arranz, L. Multiple faces of succinate beyond metabolism in blood. Haematologica 103, 1586–1592 (2018).

23. Gao, X., Chen, J., Dai, X., Zhang, D. & Zhao, Y. An Effective Strategy for Reliably Isolating Heritable and Cas9-Free Arabidopsis Mutants Generated by CRISPR/Cas9-Mediated Genome Editing. Plant Physiol 171, 1794–1800 (2016).

24. Kumar, D., Yusuf, M. A., Singh, P., Sardar, M. & Sarin, N. B. Histochemical Detection of Superoxide and H2O2 Accumulation in Brassica juncea Seedlings. Bioprotocol 4, e1108–e1108 (2014).

25. Armstrong, N. et al. SDHB knockout and succinate accumulation are insufficient for tumorigenesis but dual SDHB/NF1 loss yields SDHx-like pheochromocytomas. Cell Reports 38, 110453 (2022).

26. University, © Stanford, Stanford & California 94305. Insights into metabolism and signaling at small scale. https://purl.stanford.edu/dx038hq9888.

27. Eberlin, L. S. et al. Nondestructive, Histologically Compatible Tissue Imaging by Desorption Electrospray Ionization Mass Spectrometry. ChemBioChem 12, 2129–2132 (2011).

28. Bokhart, M. T., Nazari, M., Garrard, K. P. & Muddiman, D. C. MSiReader v1.0: Evolving Open-Source Mass Spectrometry Imaging Software for Targeted and Untargeted Analyses. J Am Soc Mass Spectrom 29, 8–16 (2018).

29. Robichaud, G., Garrard, K. P., Barry, J. A. & Muddiman, D. C. MSiReader: an open-source interface to view and analyze high resolving power MS imaging files on Matlab platform. J Am Soc Mass Spectrom 24, 718–721 (2013).

30. Schindelin, J. et al. Fiji: an open-source platform for biological-image analysis. Nat Methods 9, 676–682 (2012).

31. Laskin, J., Heath, B. S., Roach, P. J., Cazares, L. & Semmes, O. J. Tissue Imaging Using Nanospray Desorption Electrospray Ionization Mass Spectrometry. Anal. Chem. 84, 141–148 (2012).

32. Luxembourg, S. L., Mize, T. H., McDonnell, L. A. & Heeren, R. M. A. High-spatial resolution mass spectrometric imaging of peptide and protein distributions on a surface. Anal Chem 76, 5339–5344 (2004).

33. Yin, R., Burnum-Johnson, K. E., Sun, X., Dey, S. K. & Laskin, J. High spatial resolution imaging of biological tissues using nanospray desorption electrospray ionization mass spectrometry. Nat Protoc 14, 3445–3470 (2019).

34. Smith, C. A. et al. METLIN: a metabolite mass spectral database. Ther Drug Monit 27, 747–751 (2005).

35. Guijas, C. et al. METLIN: A Technology Platform for Identifying Knowns and Unknowns. Anal. Chem. 90, 3156–3164 (2018).

36. Fahy, E., Sud, M., Cotter, D. & Subramaniam, S. LIPID MAPS online tools for lipid research. Nucleic Acids Res 35, W606–W612 (2007).

37. Kim, S. et al. PubChem in 2021: new data content and improved web interfaces. Nucleic Acids Res 49, D1388–D1395 (2021).

38. Bonnington, L. S., Barcelò, D. & Knepper, T. P. Utilisation of electrospray time-of-flight mass spectrometry for solving complex fragmentation patterns: application to benzoxazinone derivatives. J Mass Spectrom 38, 1054–1066 (2003).

