## Extended Figures for "Chemical Imaging Reveals Diverse Functions of Tricarboxylic Acid Metabolites in Root Growth and Development"

### Extended Data Figures

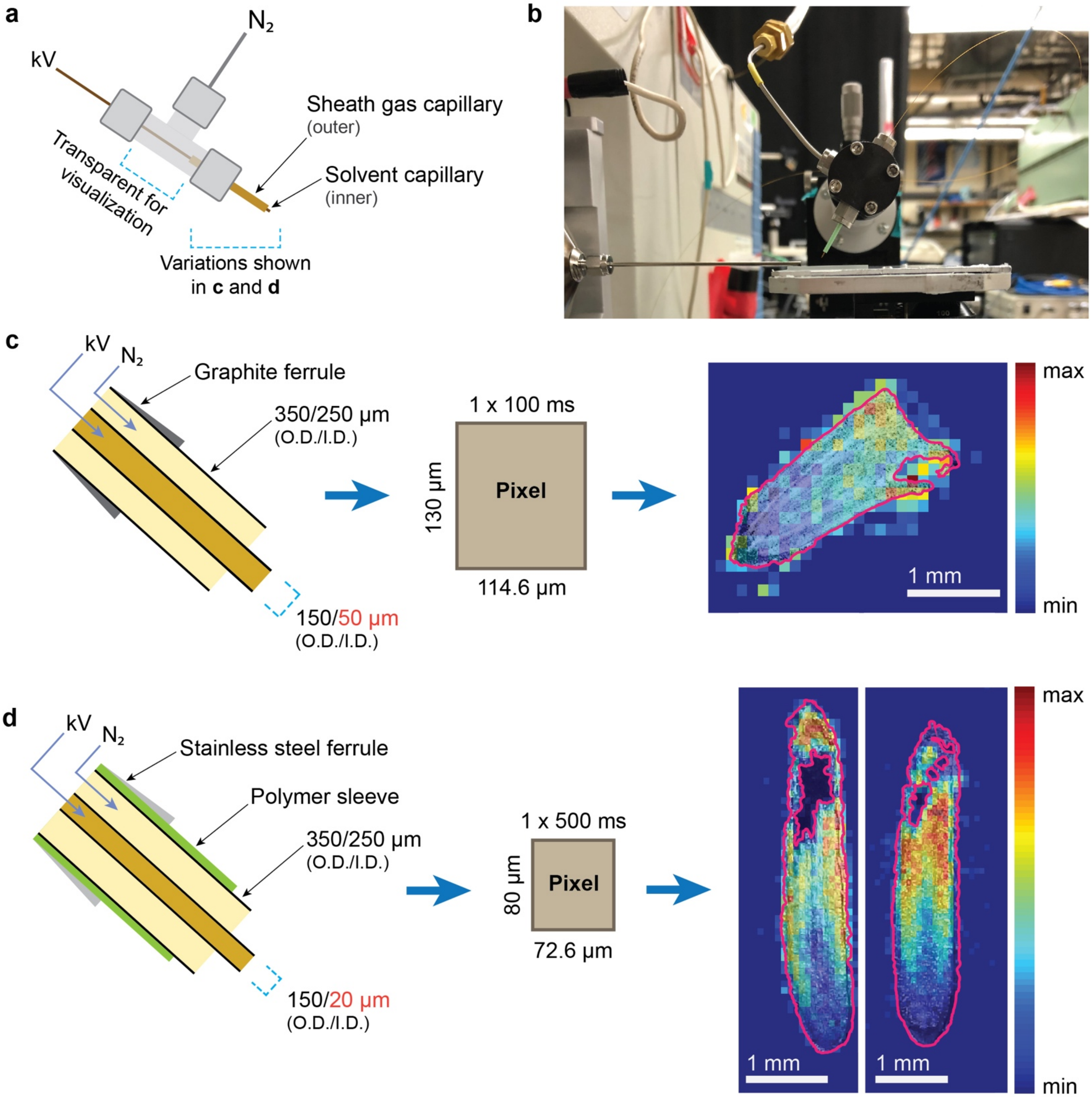


**Extended Data Fig. 1 | DESI-MSI probe design and imagine parameters for improved spatial resolution. a,** Generic schematic of the DESI probe shows the coaxial capillary construction at the tip, where the outer capillary is for the nitrogen sheath gas and the inner capillary runs continuously from the spray tip backward to the solvent syringe. **b,** Photograph of the DESI imaging set-up for maize root acquisition. Details of the coaxial probe tip, pixel size, ion injection time, and sample DESI-MS images of fumarate are shown for the initial maize imaging attempts **(c)** and the modified probe and parameters used for all image acquisition in this study **(d)**. MS images are of fumarate with brightfield microscopy image overlay. The outer borders and holes of the brightfield image are outlined in pink to demonstrate the improved resolution with the modified set-up.

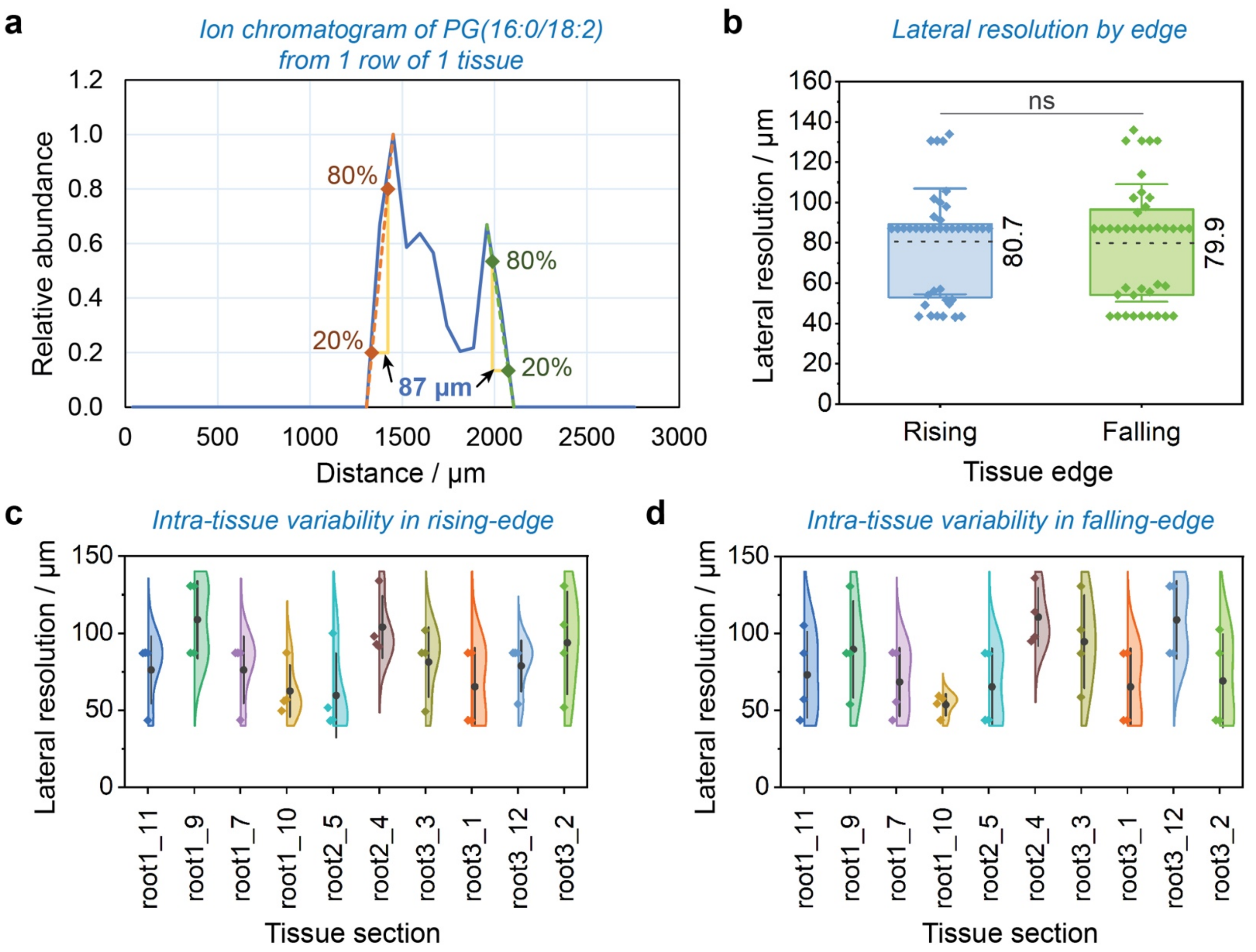


**Extended Data Fig. 2 | Lateral resolution was determined using the 80-20% rule. a,** Demonstration of the 80-20% rule using the extracted ion chromatogram of PG (16:0/18:2) from one row of one maize tissue. The distance over which the intensity rises from 20% to 80% of the maximum for each edge is the lateral resolution. **b,** Lateral resolution for the rising- and falling-edges of the ten tissue sections. Dashed lines and labels show the mean values, while whiskers are one standard deviation. The box encompasses the 25th-75th percentile. Four rows were measured for each tissue, providing n = 40 for each edge. A two-way Student's *t*-test showed no significant difference in the means at a significance level of 0.05. To justify the inclusion of four rows per tissue, we show the intra-tissue variability in the measured lateral resolution for the rising- **(c)** and falling-edge **(d)** of the ten measured tissues (n=4 per tissue). Dark gray circle and line represent the mean value and one standard deviation for each tissue section. Measured values are colored diamonds. The half-violin plots were constructed with bin widths of 20 μm.

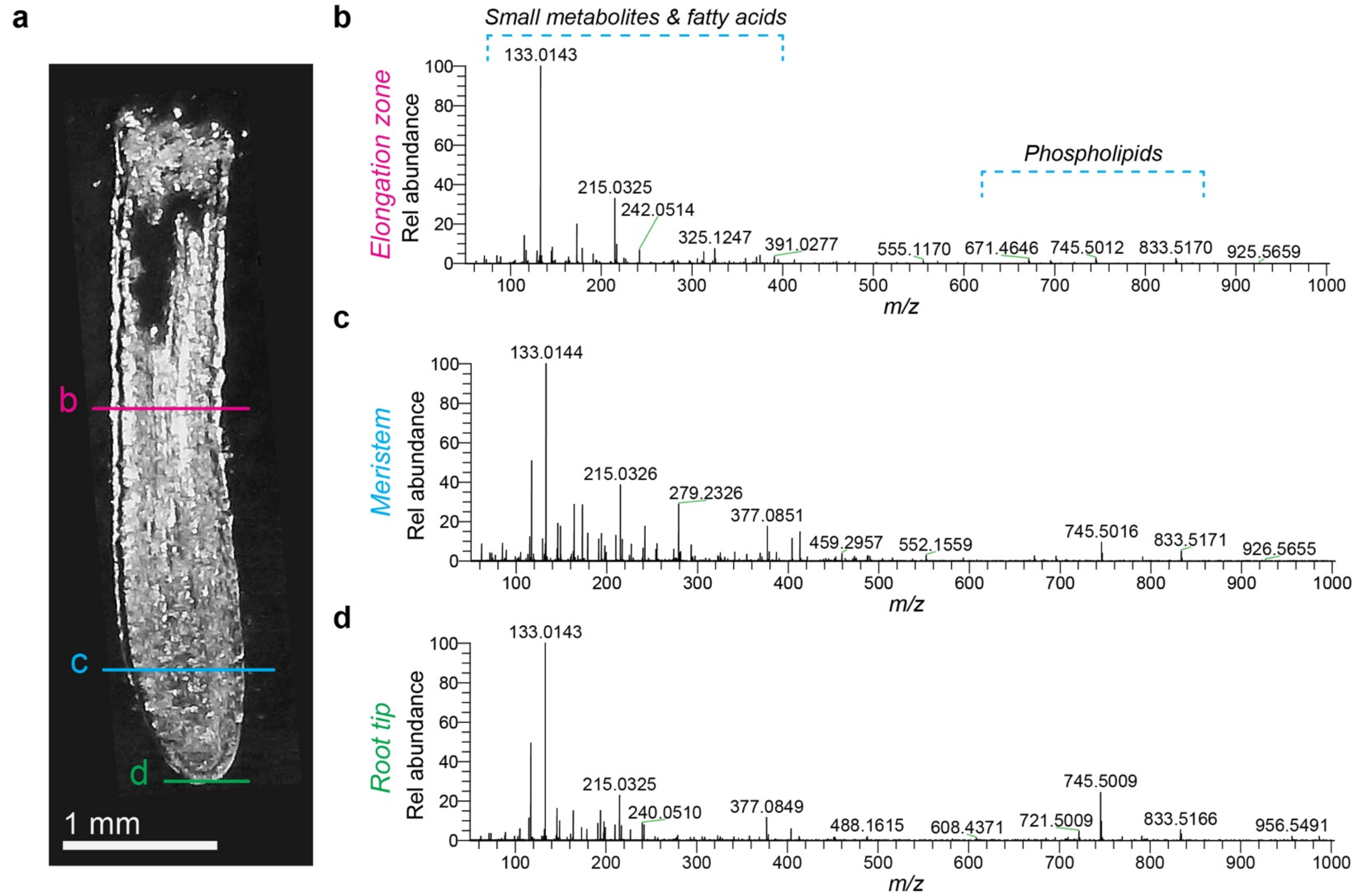


**Extended Data Fig. 3 | Representative mass spectra from distinct developmental zones of the root. a,** Brightfield microscopy image of the root section from which a representative spectrum of the elongation zone (**b**, pink), meristem zone (**c,** blue), and root tip (**d,** green) are shown. The lines in **(a)** indicate the rows that were averaged to produce the spectra in **(b-d)**.

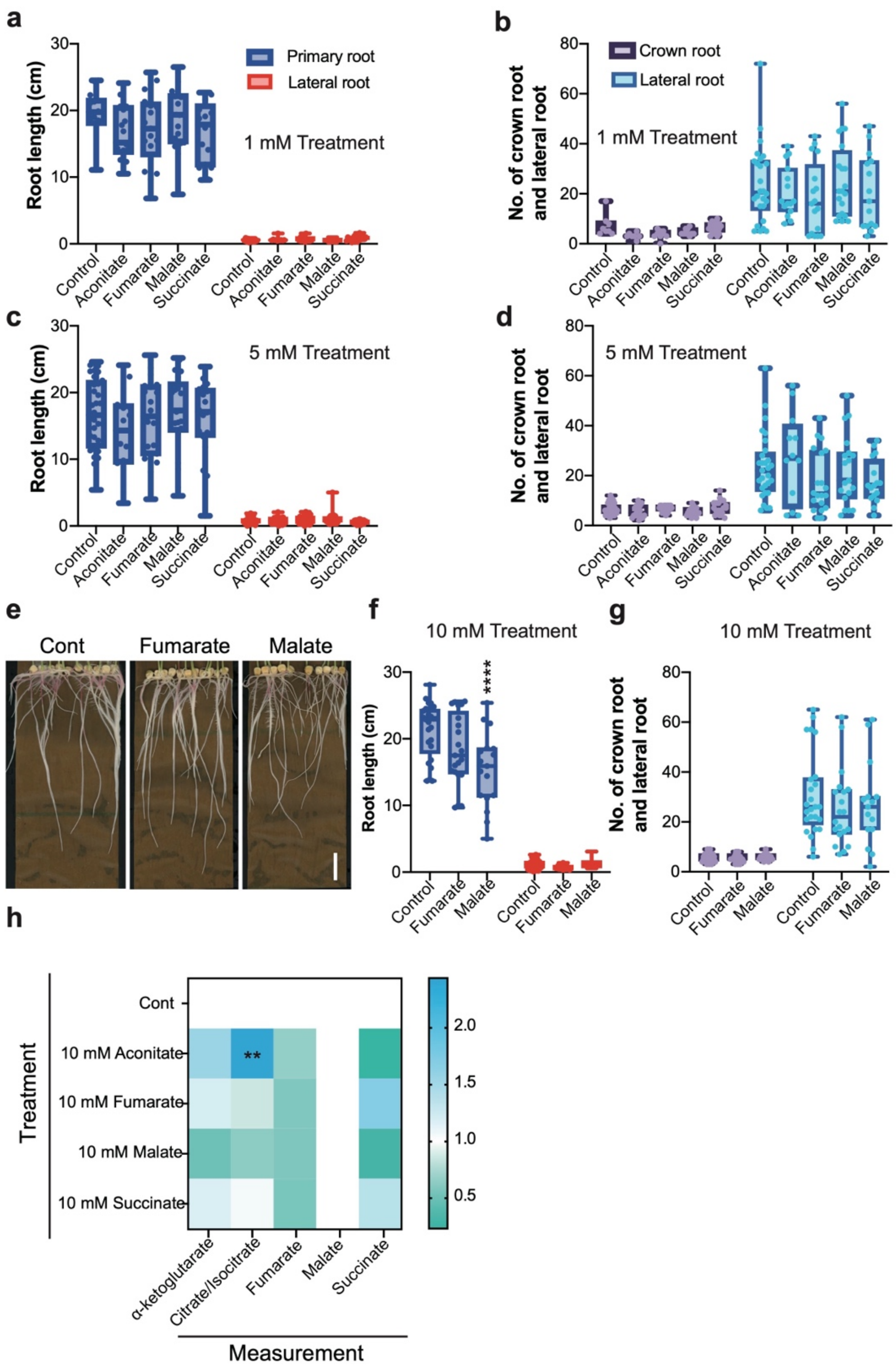


**Extended Data Fig. 4 | Maize is insensitive to TCA metabolites treatment compared to Arabidopsis. a-d,** Root phenotypes in maize seedlings treated with 1 mM and 5 mM aconitate, fumarate, malate, and succinate, respectively. **e-g,** Root phenotypes in maize seedlings treated with 10 mM fumarate, malate, and succinate. Scale bar, 3 cm. h**,** Analysis of TCA metabolites using HPLC-MS shows the effect of 10 mM metabolite treatments on the normalized level of five TCA metabolites. Data represent means ± s.d. (n =3 biological replicates). Asterisks indicate statistical significance by one-way ANOVA (**p < 0.01, ****p < 0.0001). Source data are provided as a Source Data file. Data are presented as boxplots with each dot representing the datapoint of one biological replicate. For the boxplots, the central line indicates the median, the bounds of the box show the 25th and 75th percentiles and the whiskers indicate 1.5× interquartile range.

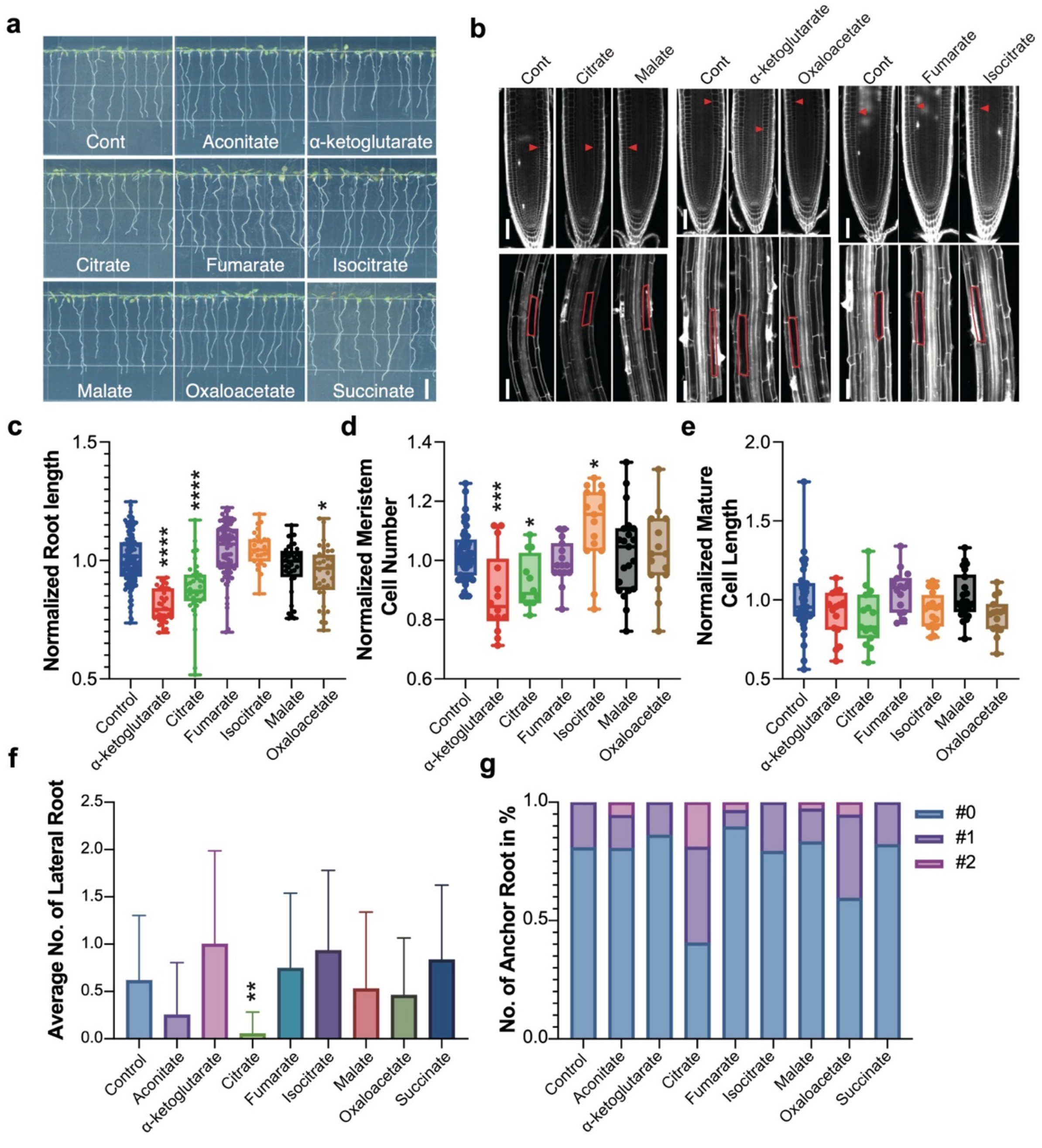


**Extended Data Fig. 5 | Response of Arabidopsis root to TCA metabolite treatments. a,** Exogenous 1 mM TCA metabolite treatments showed a different affection on plant root development. Scale bar, 1 cm. **b,** Confocal images of meristem and differentiation zone cells, respectively, under 1 mM TCA metabolite treatments. Scale bar, 50 µm. **c,** Normalized primary root length of 1 mM TCA metabolite treatments. **d,** Number of cells in the root meristem zone (MZ) after 1 mM TCA metabolite treatments. **e,** Average cell length of cells in the differentiation zone (DZ) after 1 mM TCA metabolite treatments. Data are presented as boxplots with each dot representing the datapoint of one biological replicate.

For the boxplots, the central line indicates the median, the bounds of the box show the 25th and 75th percentiles and the whiskers indicate 1.5× interquartile range. Asterisks indicate statistical significance by one-way ANOVA (*$p < 0.05$, ***$p < 0.001$, ****$p < 0.0001$). **f,** Average lateral roots number (LRN) of 1 mM TCA metabolite treatments. Data represent means ± s.d. (n = 30 plants). Asterisks indicate statistical significance by one-way ANOVA (**$p < 0.01$). **g,** Anchor roots number (ARN) of 1 mM TCA metabolite treatments. Data represent means ± s.d. (n = 30 plants).

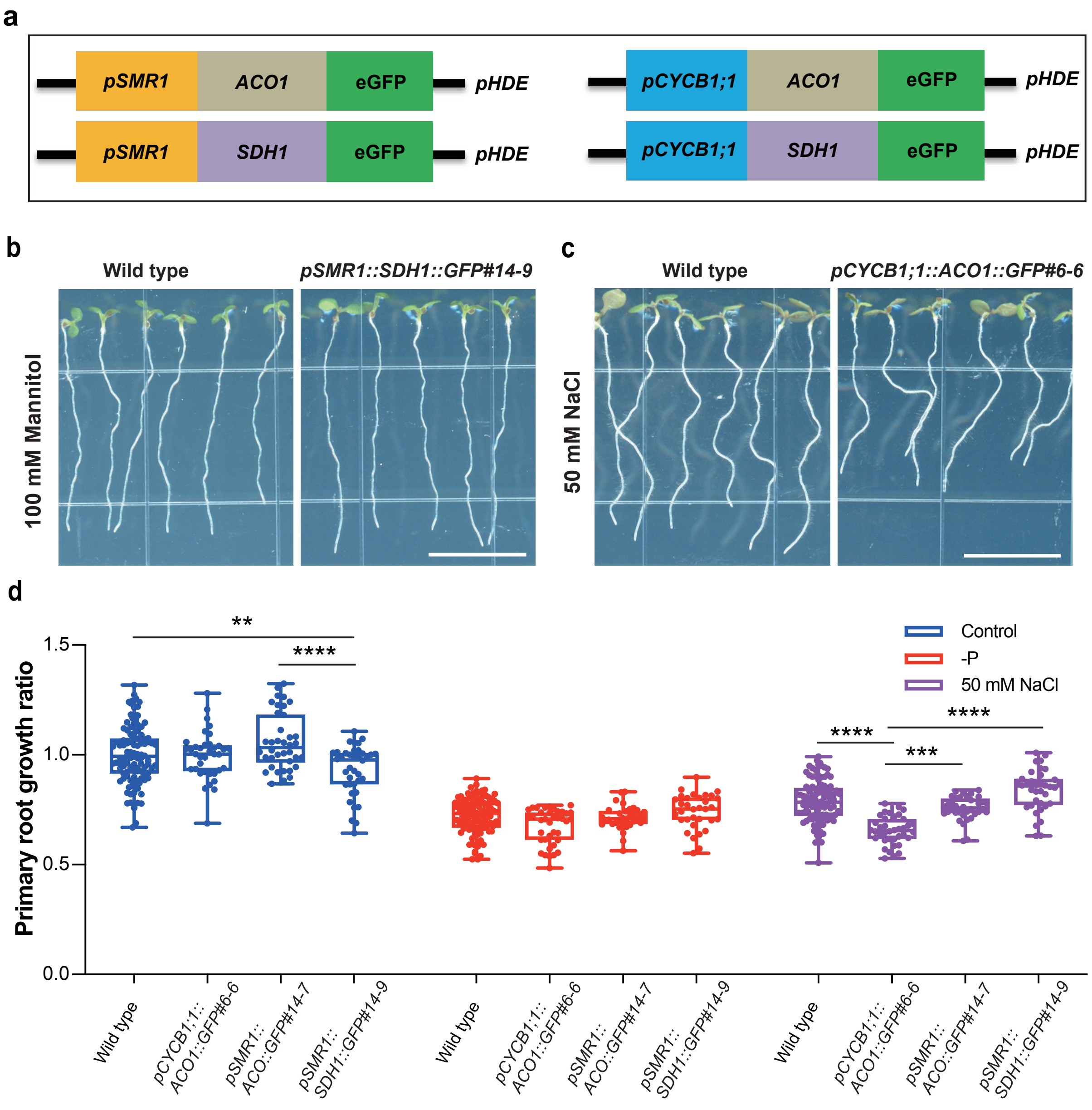


**Extended Data Fig. 6 | Ectopic expression of TCA metabolites by tissue-specific genetic engineering. a,** Schematic of constructs for ectopic expression of TCA cycle genes. **b,** Transgenic lines of *pSMR1-SDH1-GFP* show increased growth in mannitol stress compared to wild type. Scale bar, 1 cm. **c,** Transgenic lines of *pCYCB1;1-ACO1-GFP* are more sensitive to salt stress compared to wild type. Scale bar, 1 cm. **d,** Normalized root length of transgenic plants in standard media, media without phosphate, and media with 50 mM NaCl. For the boxplots, the central line indicates the median, the bounds of the box show the 25th and 75th percentiles and the whiskers indicate 1.5× interquartile range. Asterisks indicate statistical significance by one-way ANOVA (**p< 0.01, ***p < 0.001, ****P < 0.0001).

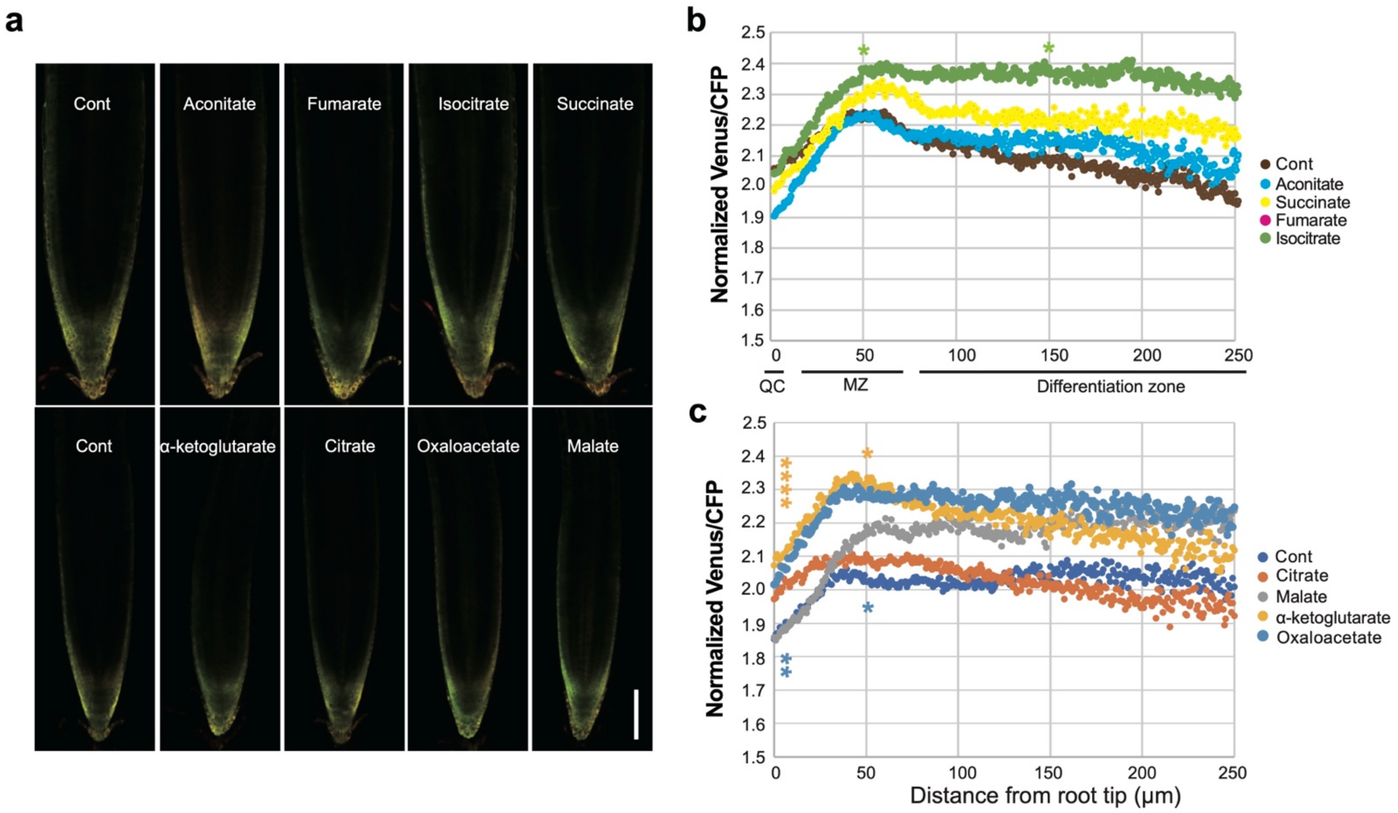


**Extended Data Fig. 7 | Response of ATP in Arabidopsis root to 1 mM TCA metabolite treatments. a,** Merged images of a confocal fluorescence (Venus and CFP) image of ATP sensor under 1 mM TCA metabolite treatments. Scale bar, 100 µm. **b-c,** Normalized Venus/CFP intensity of ATP sensor under 1 mM TCA metabolite treatments. Data represent means ± s.d. (n = 15 plants). Asterisks indicate statistical significance by one-way ANOVA (*p < 0.05, **p < 0.01, ***p < 0.001, ****p < 0.0001). Different colors indicate different TCA metabolite treatments.

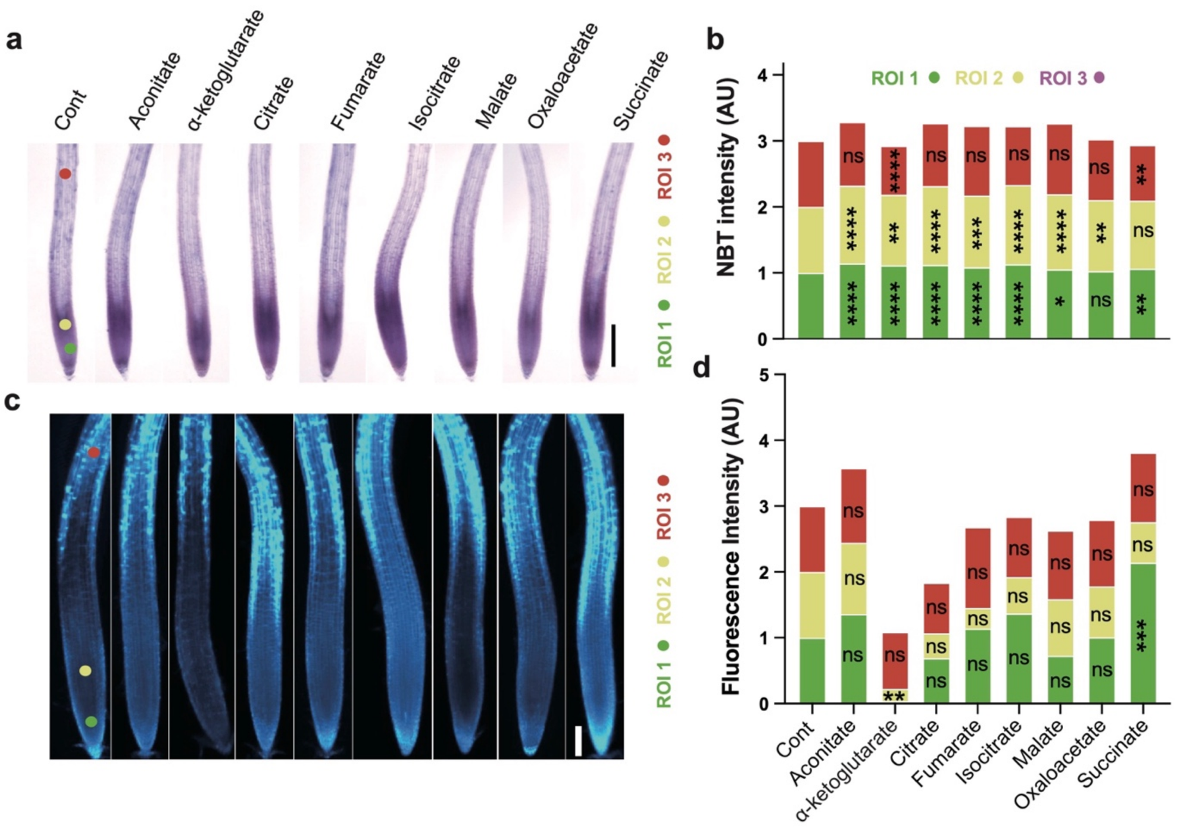


**Extended Data Figs. 8 | ROS staining of 1 mM TCA metabolite treatments of Arabidopsis. a,** Images showing NBT staining; Scale bar, 2 mm. **b,** Normalized intensity of NBT stain in three regions of interest (ROIs). **c,** Images showing $H_2DCFDA$ staining. Scale bar, 100 µm. **d,** $H_2DCFDA$ with 1 mM TCA metabolite treatments in three regions of interest (ROIs). Data represent means ± s.d. (n = 15 plants). Asterisks indicate statistical significance by one-way ANOVA (*p < 0.05, **p < 0.01, ***p < 0.001, ****p < 0.0001).

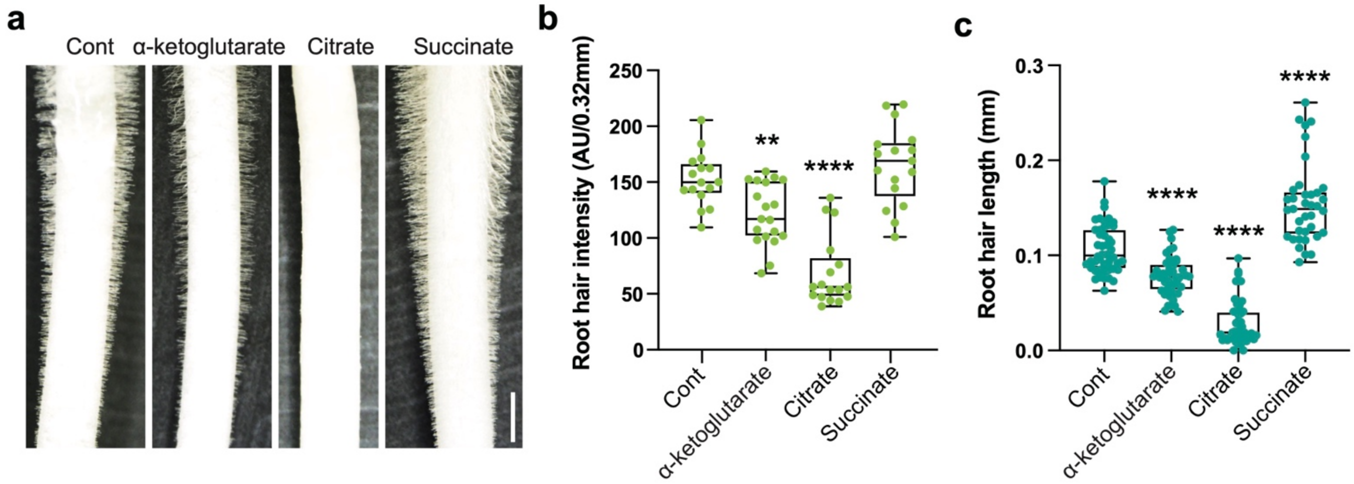


**Extended Data Fig.9 | Root hair phenotype of 10 mM TCA metabolite treatments in maize. a,** Root hair phenotype of 10 mM α-ketoglutarate, citrate, and succinate treatment in maize. Scale bar, 0.15 mm. **b,** Normalized root hair density under different treatments. **c,** Root hair length under different treatments. For the boxplots, the central line indicates the median, the bounds of the box show the 25th and 75th percentiles and the whiskers indicate 1.5× interquartile range. Asterisks indicate statistical significance by one-way ANOVA (**$p < 0.01$, ****$p < 0.0001$).
