## Supplementary Information for "Chemical Imaging Reveals Diverse Functions of Tricarboxylic Acid Metabolites in Root Growth and Development"

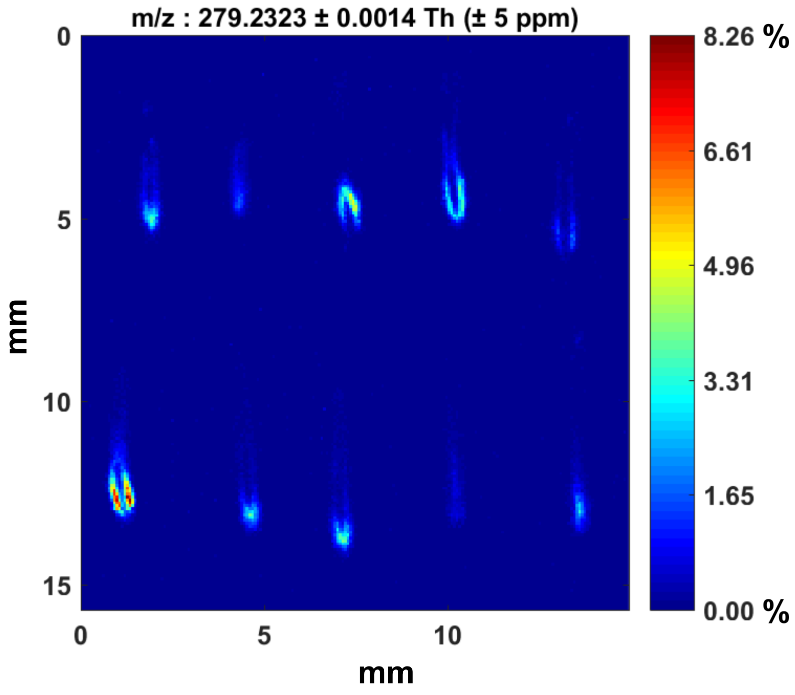


**Supplementary Fig. 1 |** DESI-MS images of fatty acid(18:2) (*m/z* 279.2323) from all imaged B73 maize root sections (across 3 biological replicates). Images are normalized to the total ion current, and the maximum intensity is 8.263%.

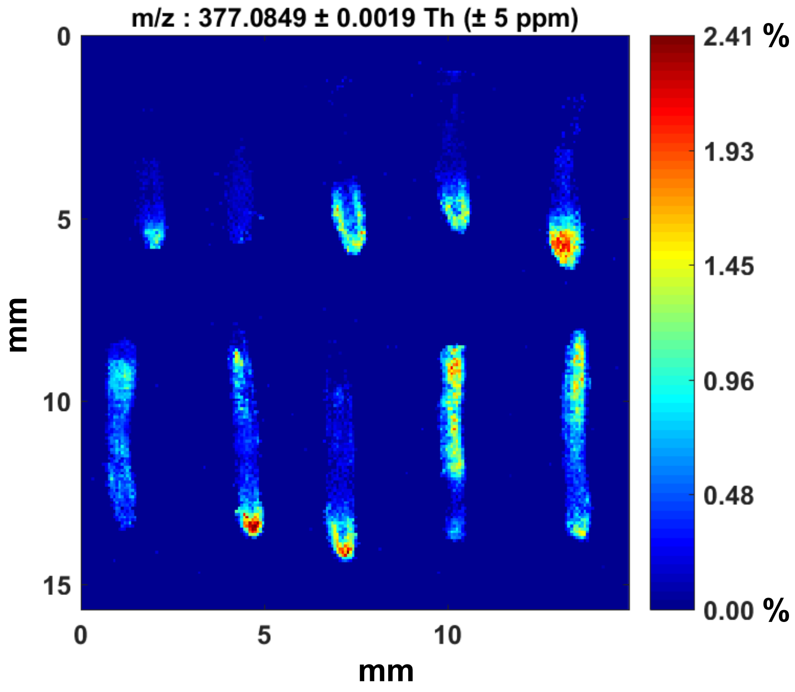


**Supplementary Fig. 2 |** DESI-MS images of hexose-hexose (*m/z* 377.0849) from all imaged B73 maize root sections (across 3 biological replicates). Images are normalized to the total ion current, and the maximum intensity is 2.412%.

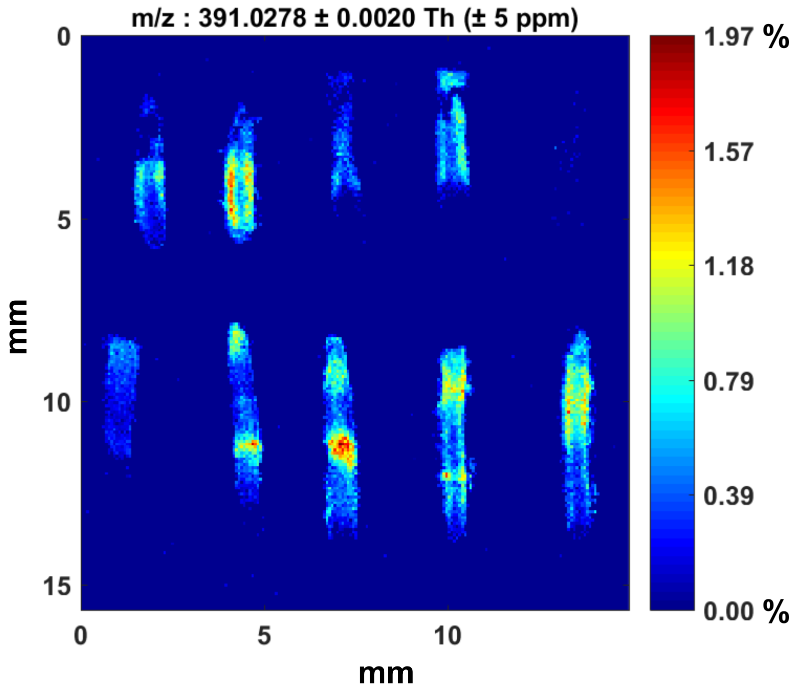


**Supplementary Fig. 3 |** DESI-MS images of unknown 1 (*m/z* 391.0278) from all imaged B73 maize root sections (across 3 biological replicates). Images are normalized to the total ion current, and the maximum intensity is 1.966%.

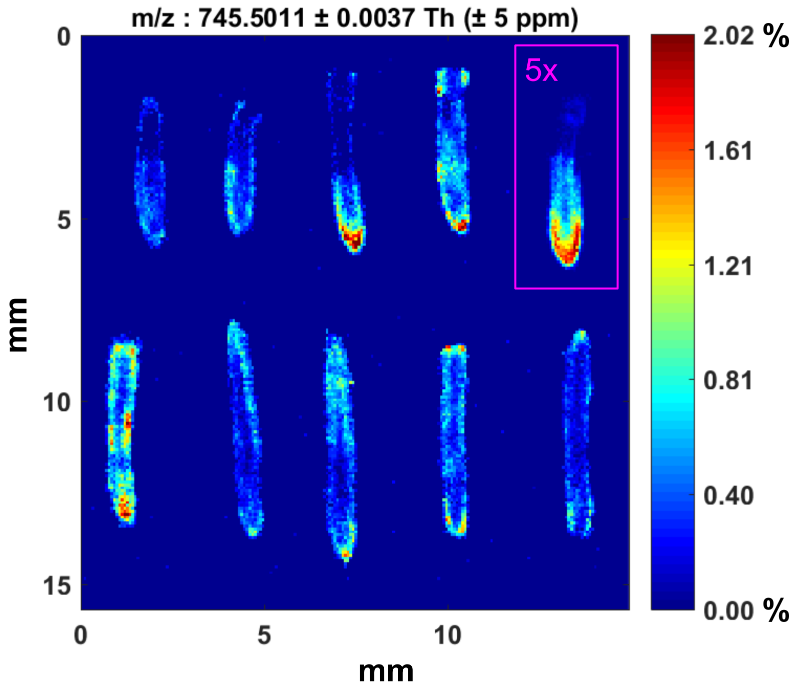


**Supplementary Fig. 4 |** DESI-MS images of phosphatidylglycerol (16:0/18:2) (*m/z* 745.5011) from all imaged B73 maize root sections (across 3 biological replicates). Images are normalized to the total ion current, and the maximum intensity is 2.016%, except for the root section outlined in pink. For that section only, the max intensity is 5x higher: 10.078%.

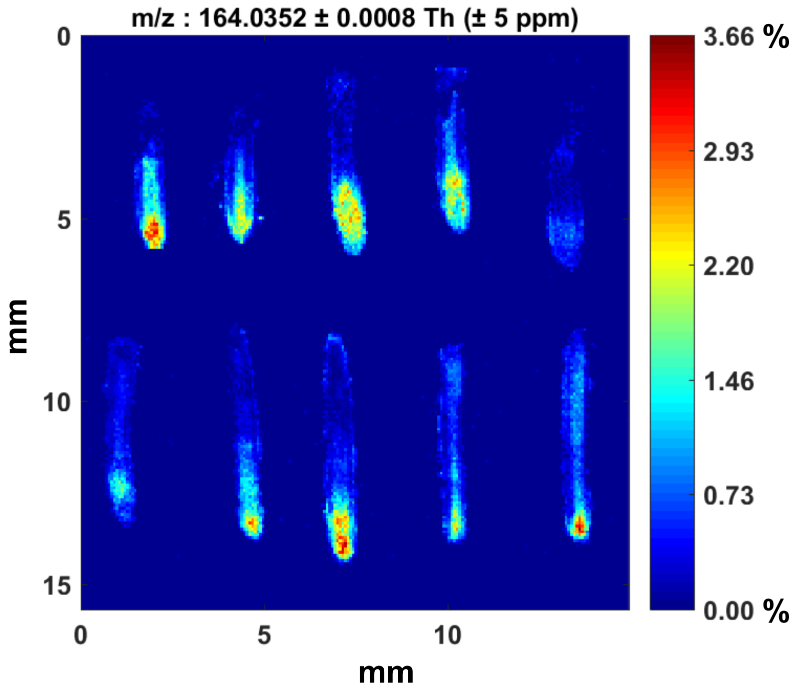


**Supplementary Fig. 5 |** DESI-MS images of MBOA (*m/z* 164.0352) from all imaged B73 maize root sections (across 3 biological replicates). Images are normalized to the total ion current, and the maximum intensity is 3.662%.

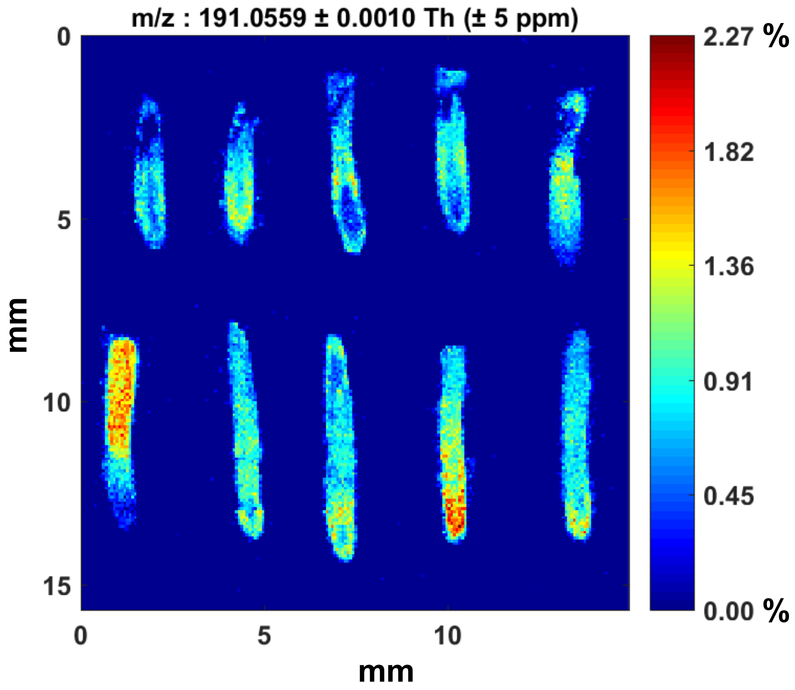


**Supplementary Fig. 6 |** DESI-MS images of quinate (*m/z* 191.0559) from all imaged B73 maize root sections (across 3 biological replicates). Images are normalized to the total ion current, and the maximum intensity is 2.27%.

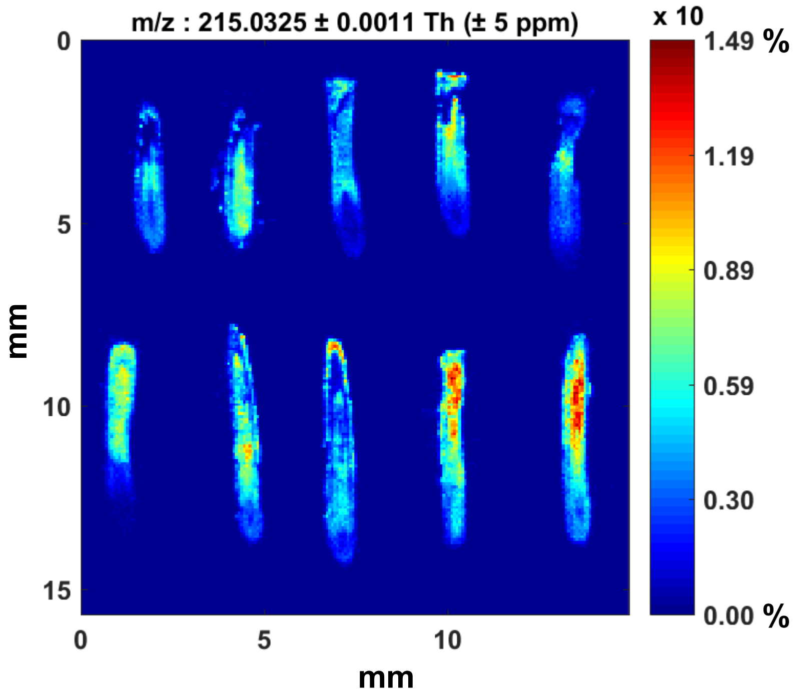


**Supplementary Fig. 7 |** DESI-MS images of hexose (*m/z* 215.0325) from all imaged B73 maize root sections (across 3 biological replicates). Images are normalized to the total ion current, and the maximum intensity is 14.853%.

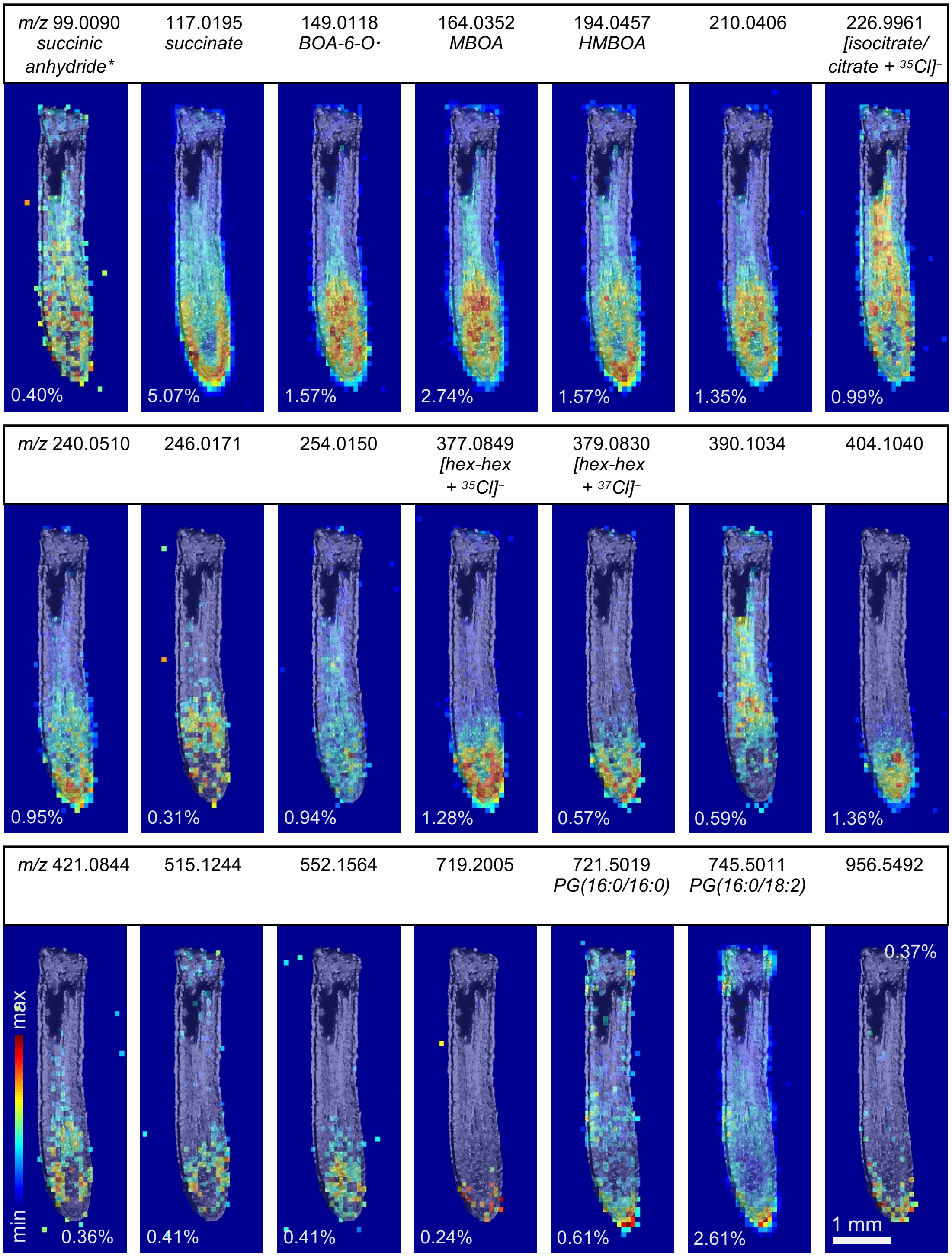


**Supplementary Fig. 8 | DESI-MS images of tip- and stele-localized compounds with brightfield overlay.** Some peaks have been identified, while others are unknown, highlighting the potential of this technique for discovery. The maximum peak intensity as a percentage of TIC is shown for each image. Identifications indicated with an asterisk are not confirmed by MS/MS, meaning the peak could be an isomer of the given compound. All other named compounds have MS/MS patterns included in the supplementary information. Ions are deprotonated unless otherwise indicated.

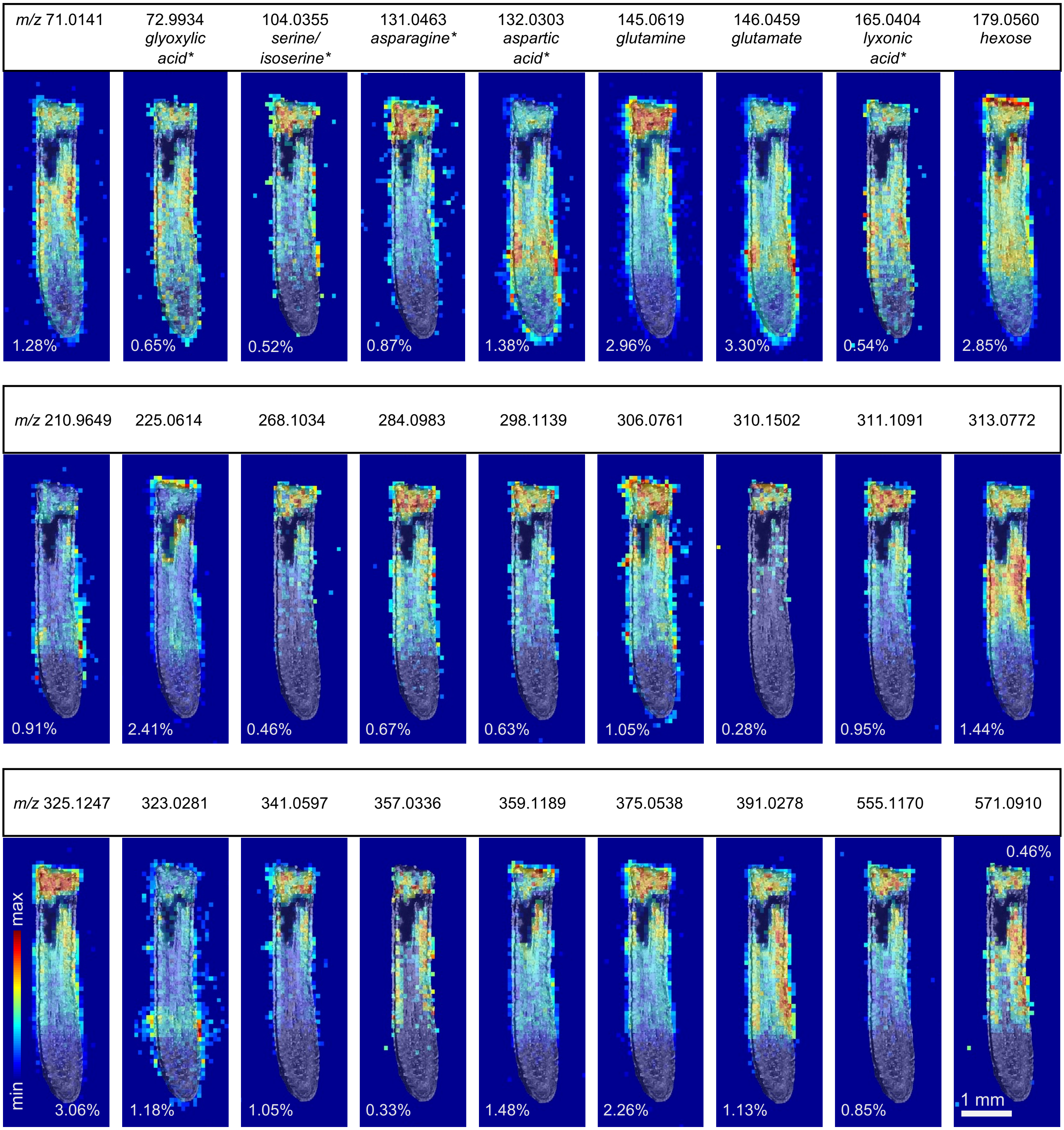


**Supplementary Fig. 9 | DESI-MS images of additional compounds enriched in the elongation and differentiation zones with brightfield overlay.** Some peaks have been identified, while others are unknown, highlighting the potential of this technique for discovery. The maximum peak intensity as a percentage of TIC is shown for each image. Identifications indicated with an asterisk are not confirmed by MS/MS, meaning the peak could be an isomer of the given compound. All other named compounds have MS/MS patterns included in the supplementary information. Ions are deprotonated unless otherwise indicated.

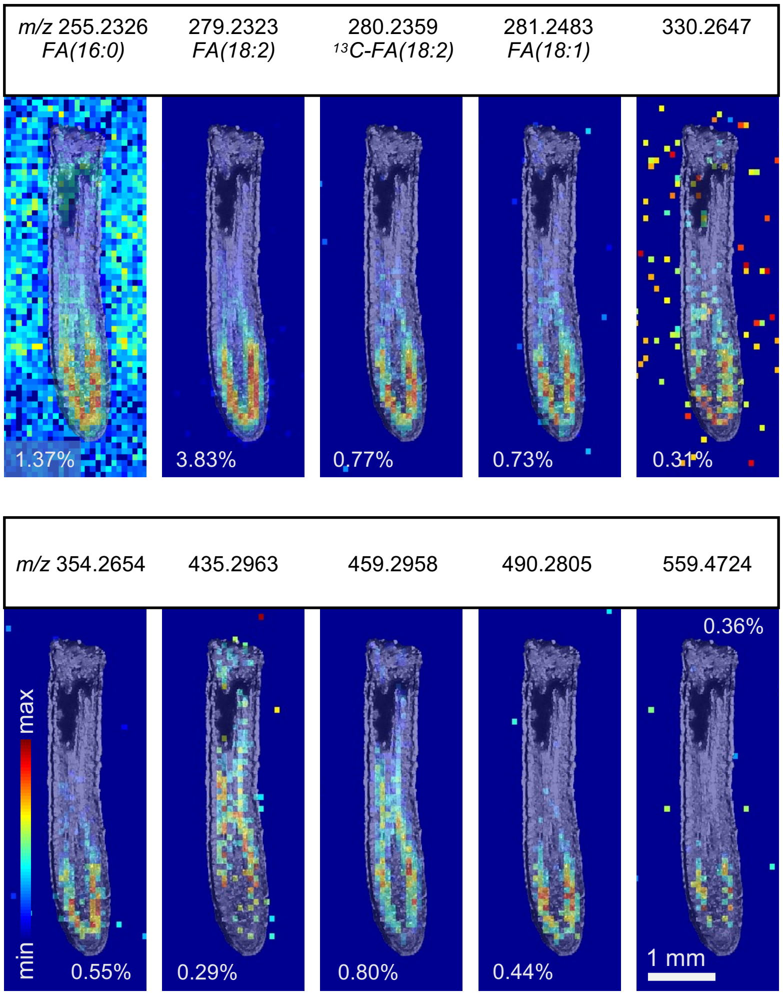


**Supplementary Fig. 10 | DESI-MS images of compounds enriched in the endodermis/cortex with brightfield overlay.** Some peaks have been identified, while others are unknown, highlighting the potential of this technique for discovery. The maximum peak intensity as a percentage of TIC is shown for each image. MS/MS patterns are included in the supplementary information for named peaks. Ions are deprotonated.

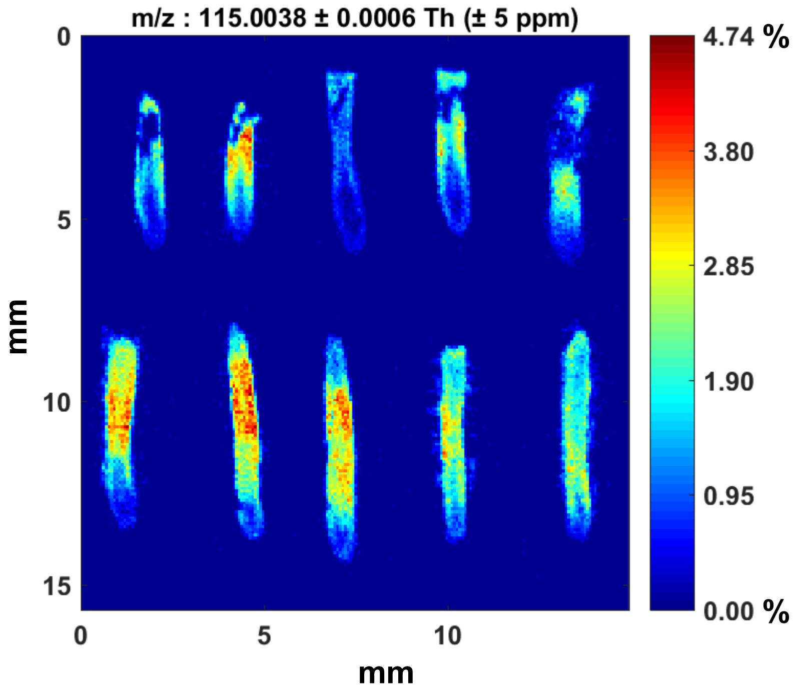


**Supplementary Fig. 11 |** DESI-MS images of fumarate (*m/z* 115.0038) from all imaged B73 maize root sections (across 3 biological replicates). Images are normalized to the total ion current, and the maximum intensity is 4.744%.

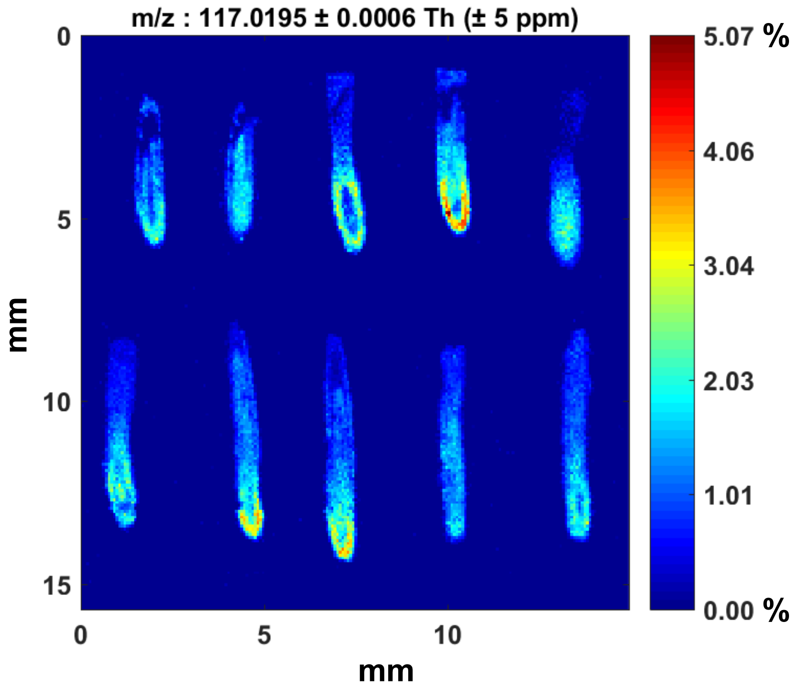


**Supplementary Fig. 12 |** DESI-MS images of succinate (*m/z* 117.0195) from all imaged B73 maize root sections (across 3 biological replicates). Images are normalized to the total ion current, and the maximum intensity is 5.07%.

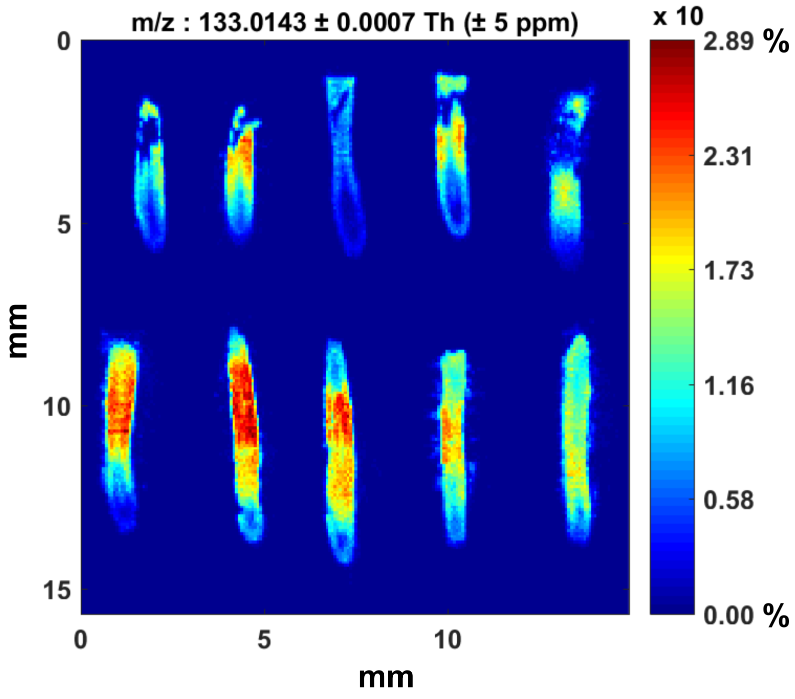


**Supplementary Fig. 13 |** DESI-MS images of malate (*m/z* 133.0143) from all imaged B73 maize root sections (across 3 biological replicates). Images are normalized to the total ion current, and the maximum intensity is 28.897%.

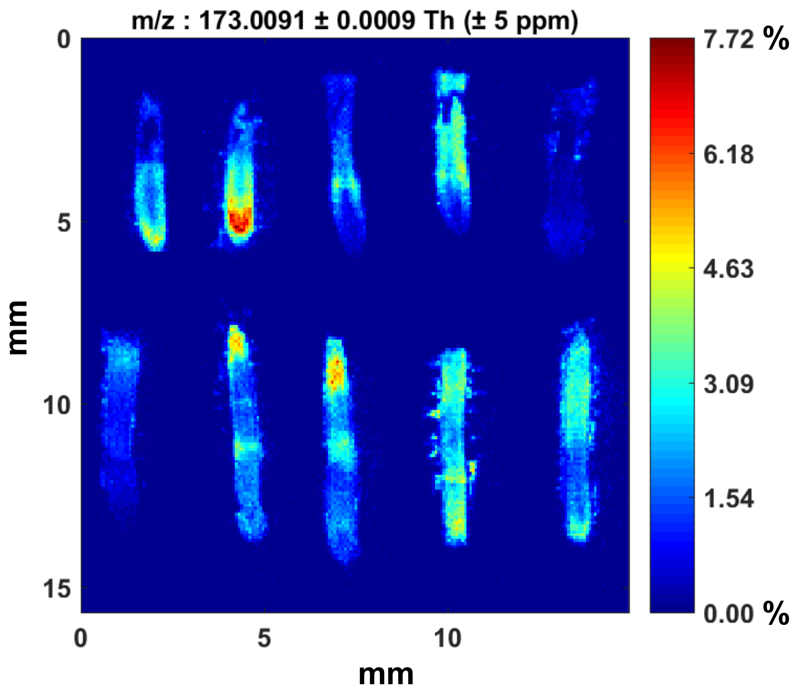


**Supplementary Fig. 14 |** DESI-MS images of aconitate (*m/z* 173.0091) from all imaged B73 maize root sections (across 3 biological replicates). Images are normalized to the total ion current, and the maximum intensity is 7.722%.

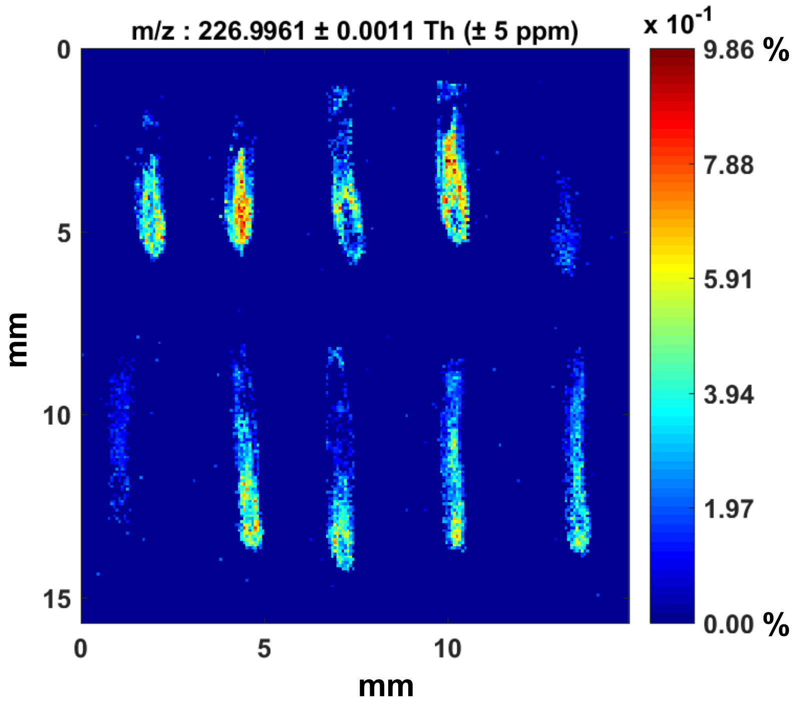


**Supplementary Fig. 15 |** DESI-MS images of citrate/isocitrate (*m/z* 226.9961) from all imaged B73 maize root sections (across 3 biological replicates). Images are normalized to the total ion current, and the maximum intensity is 0.986%.

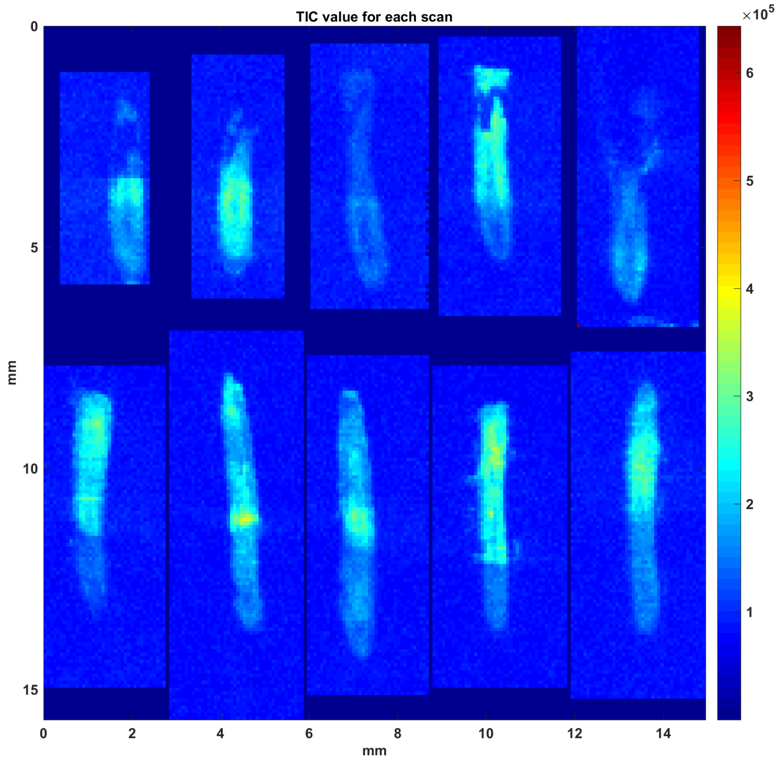


**Supplementary Fig. 16 |** MS images of the total ion current (TIC) by pixel for each of the ten imaged B73 maize root sections. The maximum intensity is 643069.

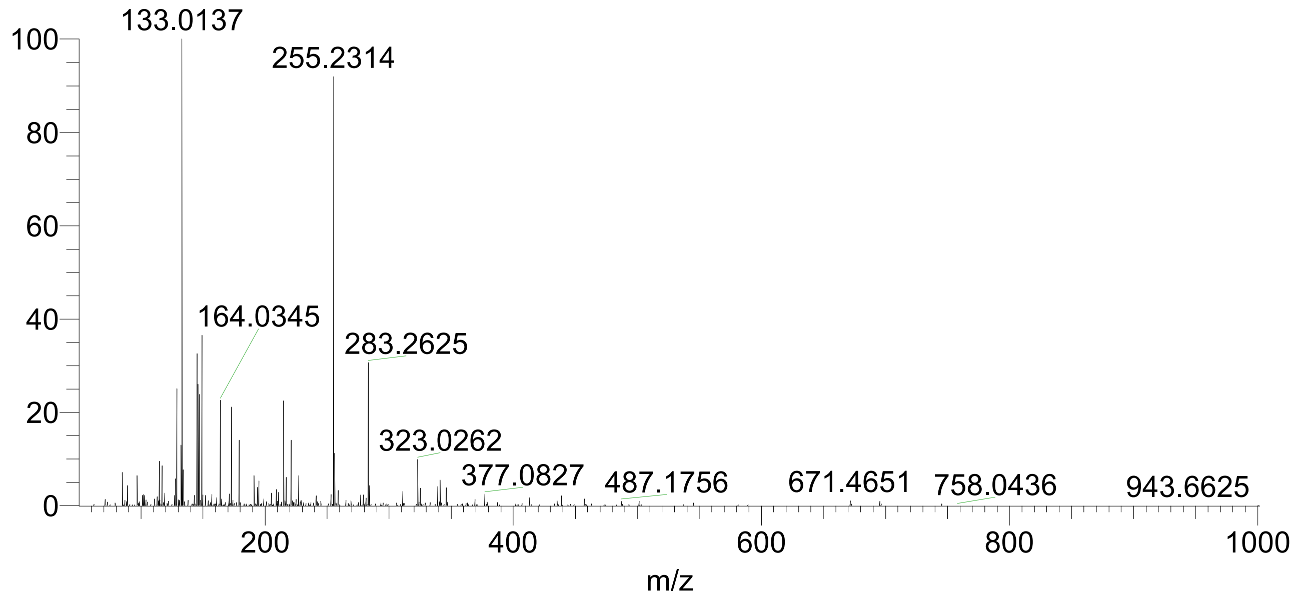


**Supplementary Fig. 17 |** Full mass spectrum of B73 maize root extract used for MS/MS analysis.

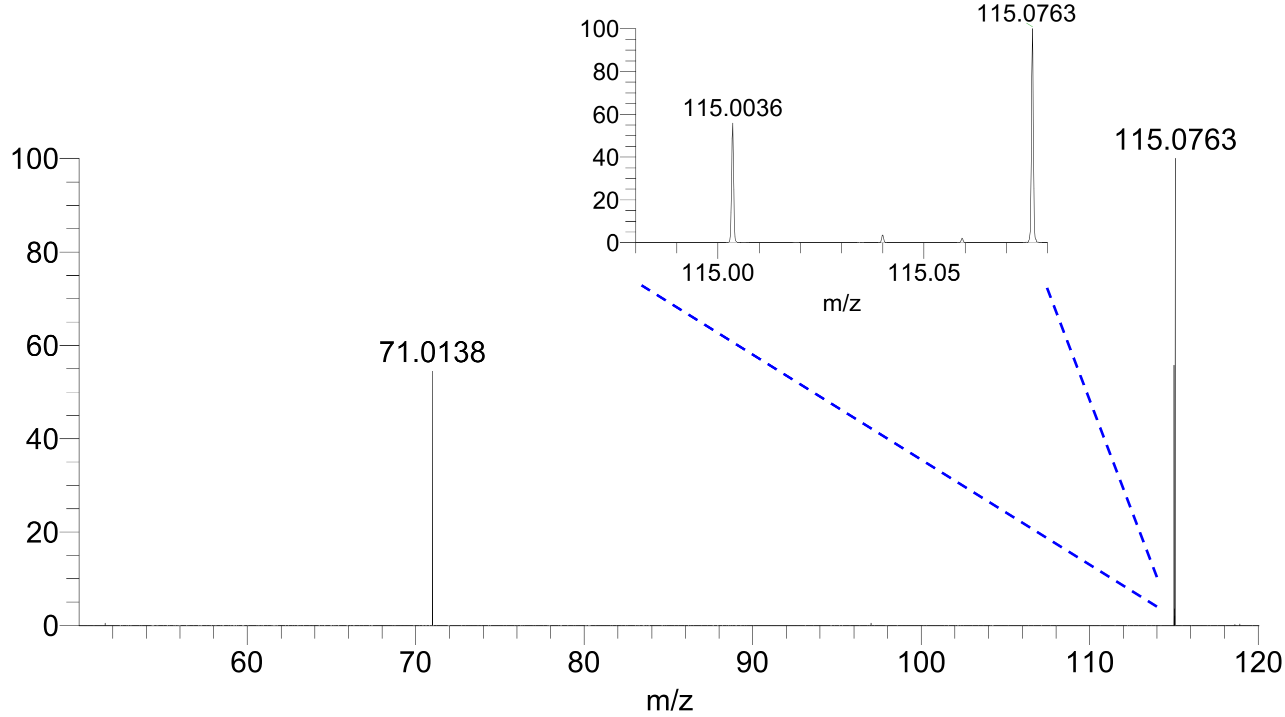


**Supplementary Fig. 18 |** MS/MS spectrum of fumarate (*m/z* 115.0036) with NCE 20%. Inset zooms in on isolated parent peak window to show fumarate.

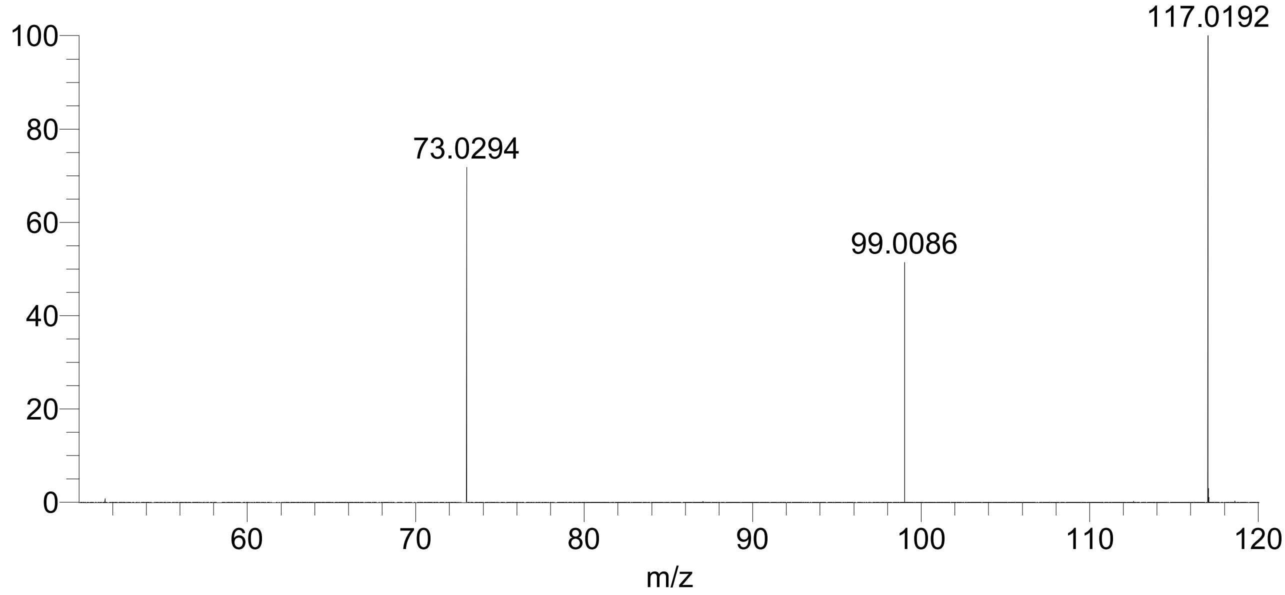


**Supplementary Fig. 19 |** MS/MS spectrum of succinate (*m/z* 117.0192) with NCE 20%.

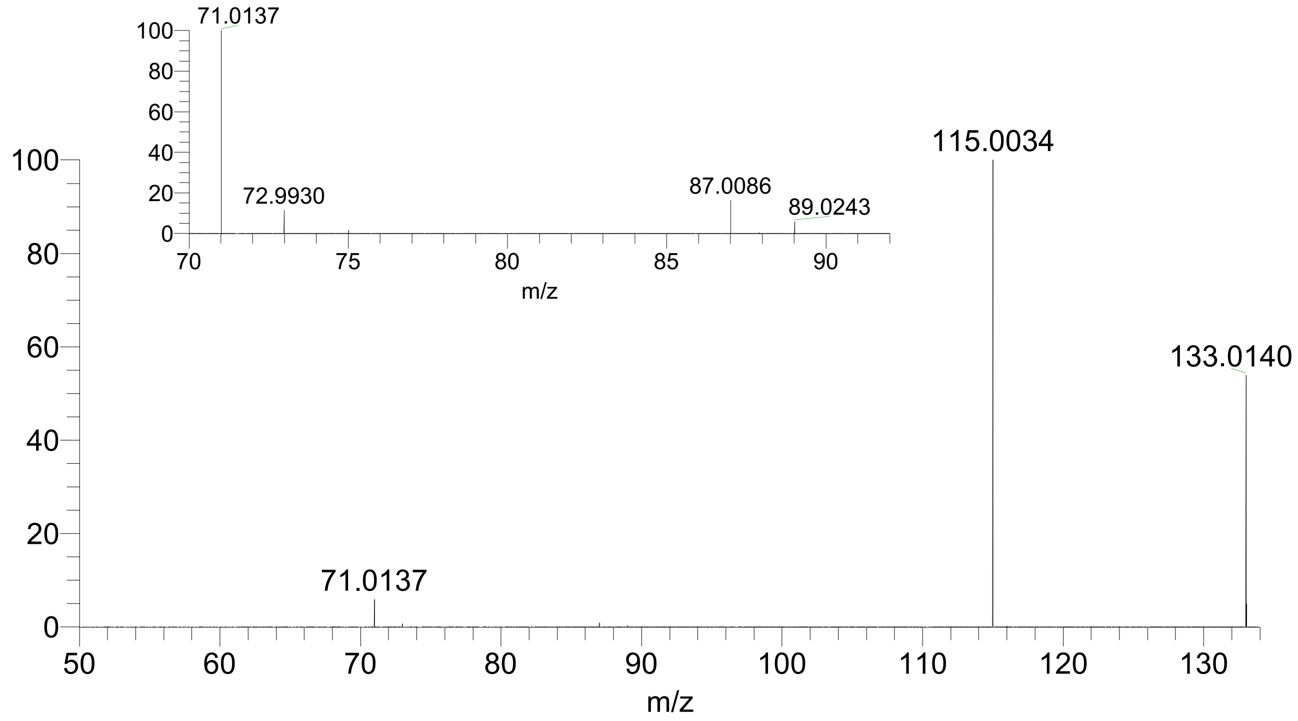


**Supplementary Fig. 20 |** MS/MS spectrum of malate (*m/z* 133.0140) with NCE 20%. Inset expands *m/z* 70-92 to help visualize low abundance fragments.

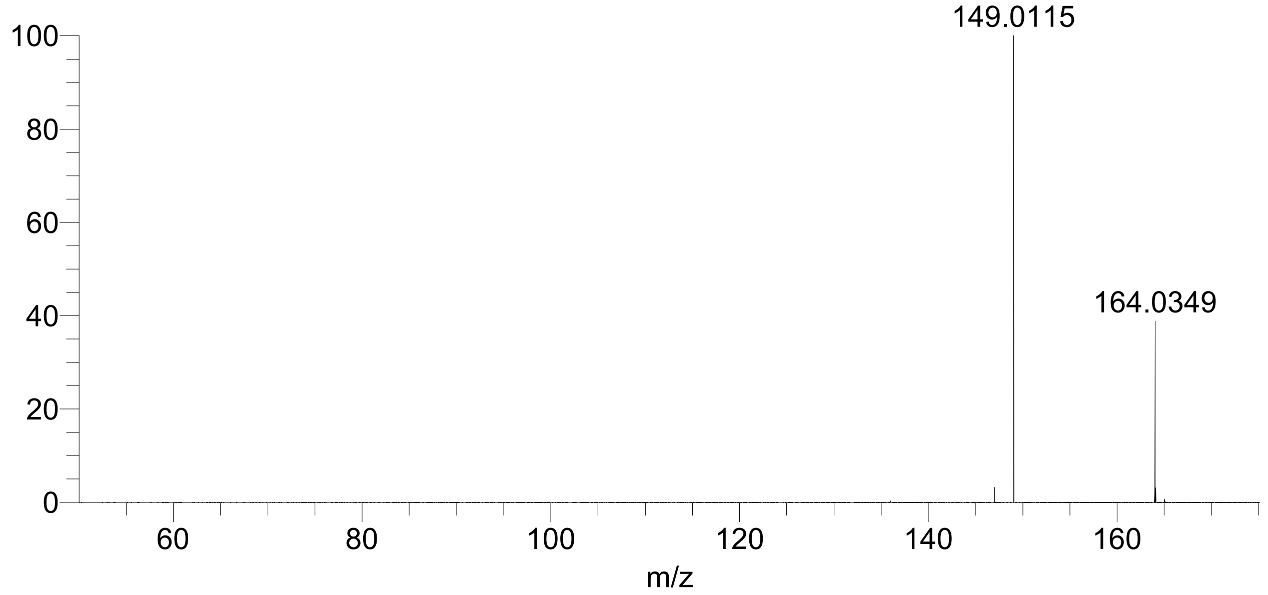


**Supplementary Fig. 21 |** MS/MS spectrum of MBOA (*m/z* 164.0349) with NCE 25%.

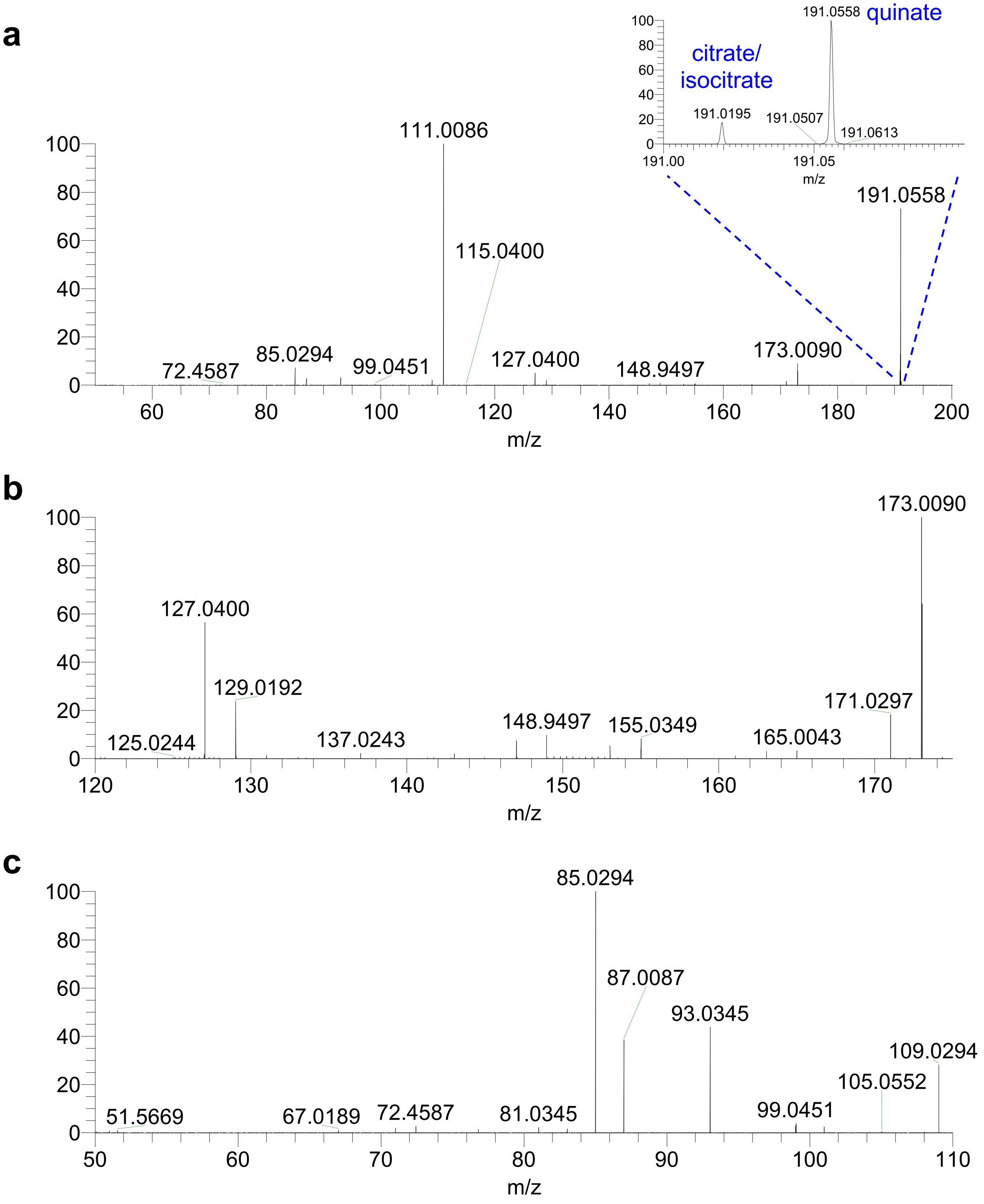


**Supplementary Fig. 22 | a,** MS/MS spectrum of quinate (*m/z* 191.0558) and citrate/isocitrate (*m/z* 191.0195) with NCE 25%. Inset expands to show both parent peaks, which could not be individually isolated. Zoom in on *m/z* 120-175 (**b**) and *m/z* 50-110 (**c**) to better visualize low abundance fragments.

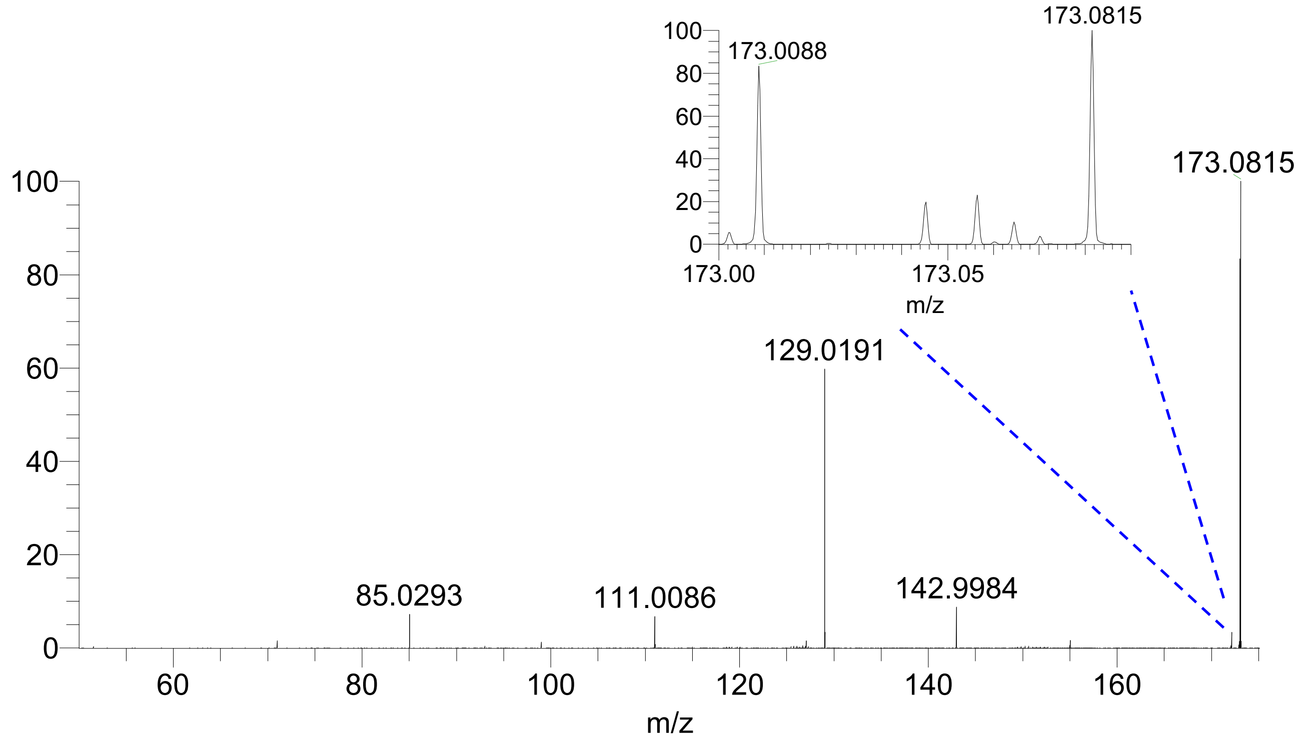


**Supplementary Fig. 23 |** MS/MS spectrum of aconitate (*m/z* 173.0088) with NCE 15%. The isolation window also captured *m/z* 173.0815, an unknown, as shown by the inset.

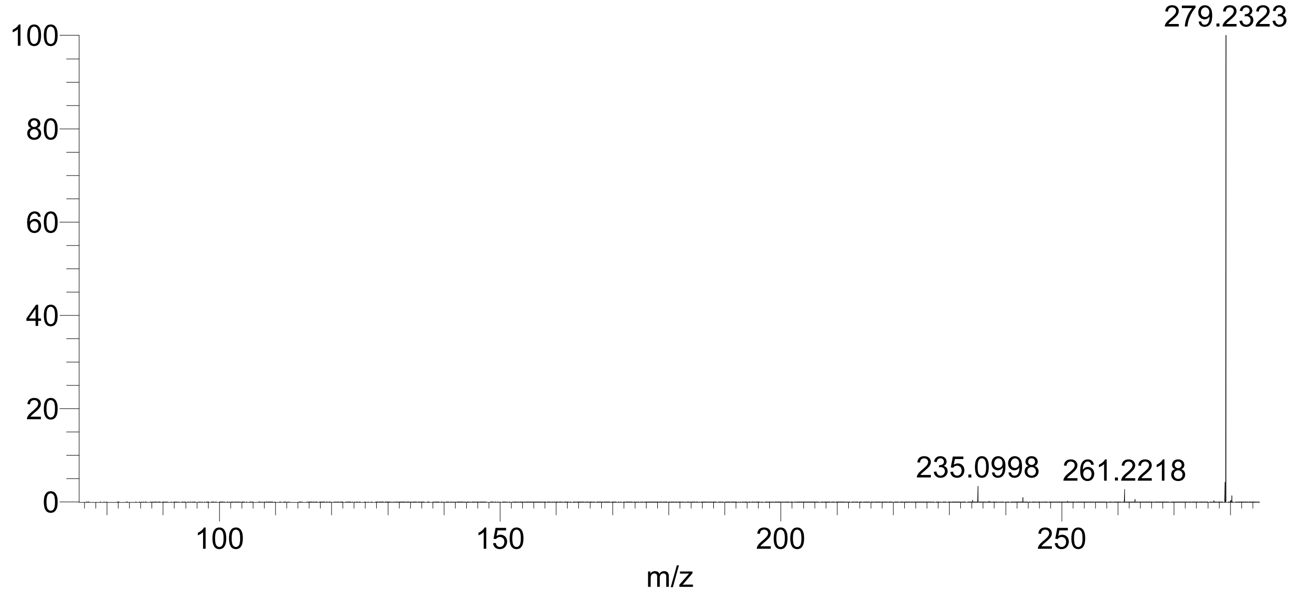


**Supplementary Fig. 24 |** MS/MS spectrum of FA(18:2) (*m/z* 279.2323) with NCE 30%.

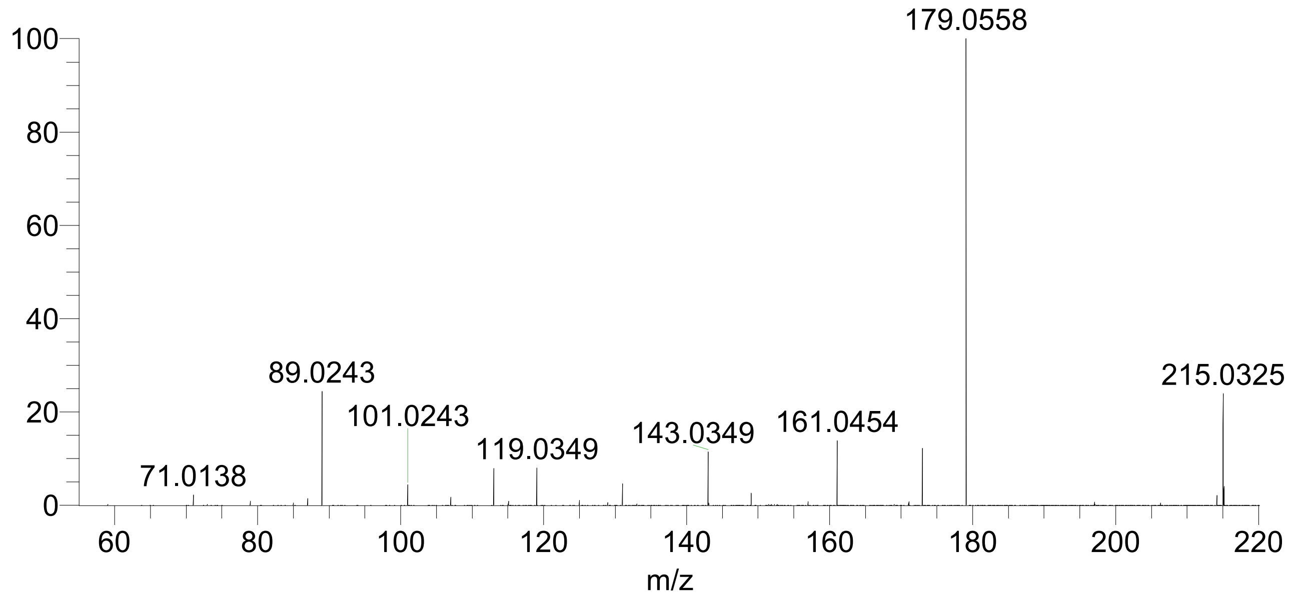


**Supplementary Fig. 25 |** MS/MS spectrum of hexose (*m/z* 215.0325) with NCE 20%. *m/z* 215.0325 represents the $[M+Cl]^-$ peak, and *m/z* 179.0558 represents the $[M-H]^-$ peak. The exact monosaccharide isomer cannot be definitively identified.

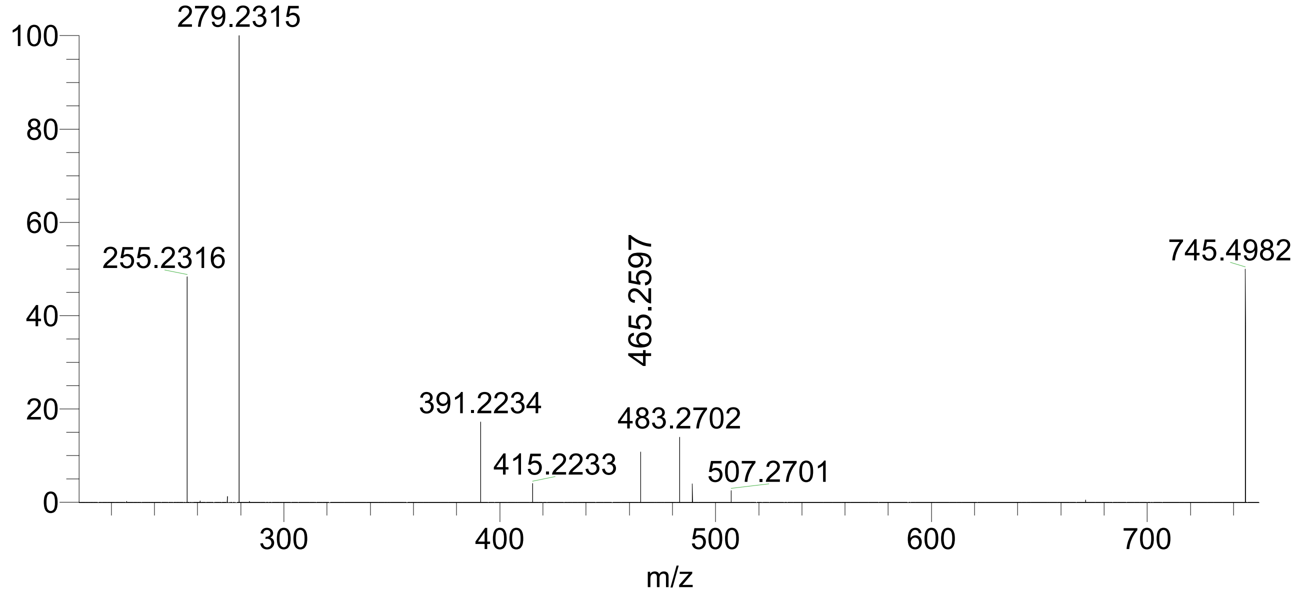


**Supplementary Fig. 26 |** MS/MS spectrum of PG(16:0/18:2) (*m/z* 745.4982) with NCE 30%.

**a**

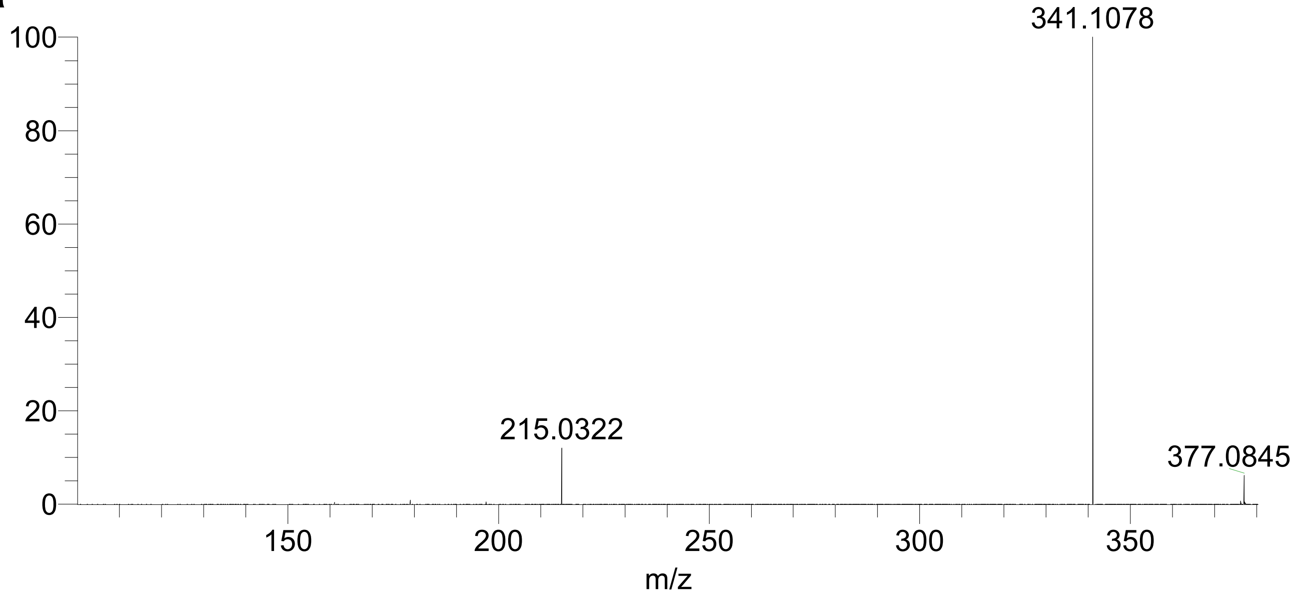


**b**

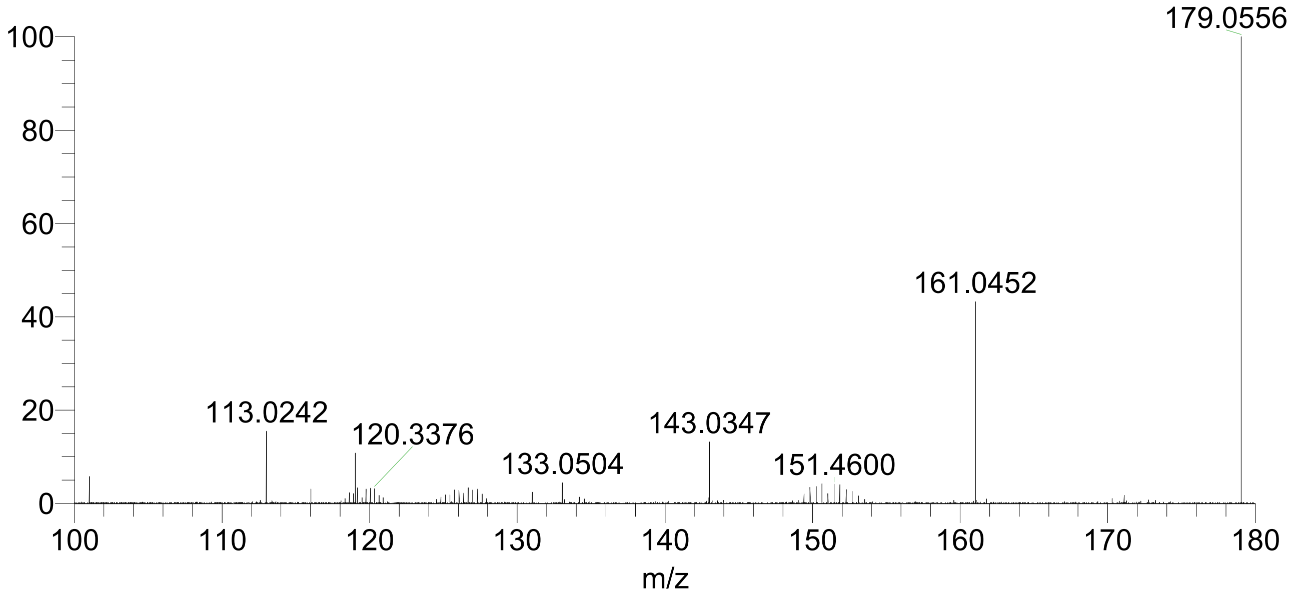


**Supplementary Fig. 27 | a,** MS/MS spectrum of hexose-hexose (*m/z* 377.0845) with NCE 22%. *m/z* 377.0845 represents the $[M+Cl]^-$ peak, and *m/z* 341.1078 represents the $[M-H]^-$ peak. **b,** Zoom in on *m/z* 100-180 better visualizes the low abundance peaks. The specific disaccharide isomer cannot be definitively identified.

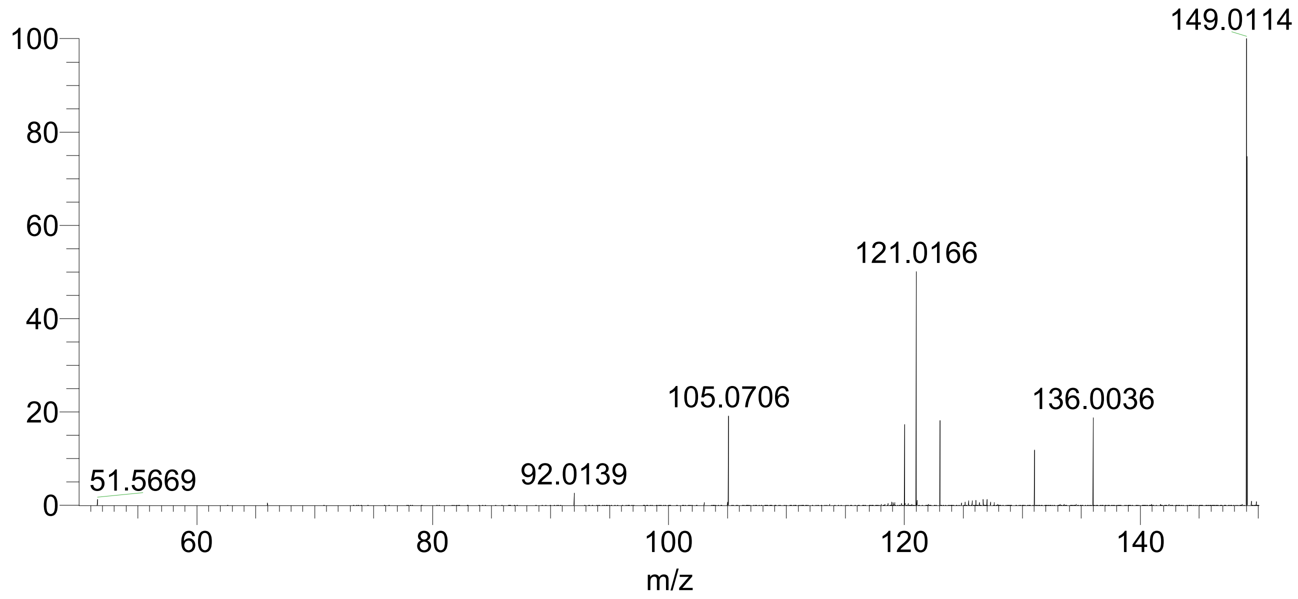


**Supplementary Fig. 28 |** MS/MS spectrum of BOA-6-O· (*m/z* 149.0114, $C_7H_3NO_3^{\cdot -}$) with NCE 25%. Expected fragment is *m/z* 121.0166.

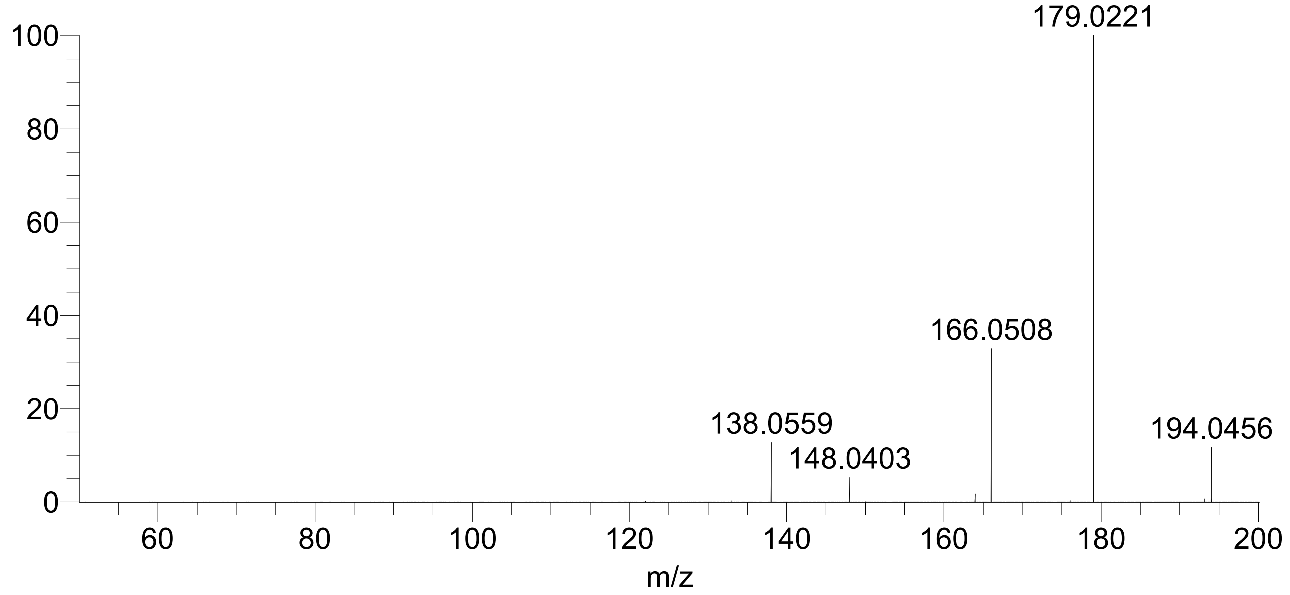


**Supplementary Fig. 29 |** MS/MS spectrum of HMBOA, 2-hydroxy-7-methoxy-1,4-benzoxazine-3-one, (*m/z* 194.0456) with NCE 25%.

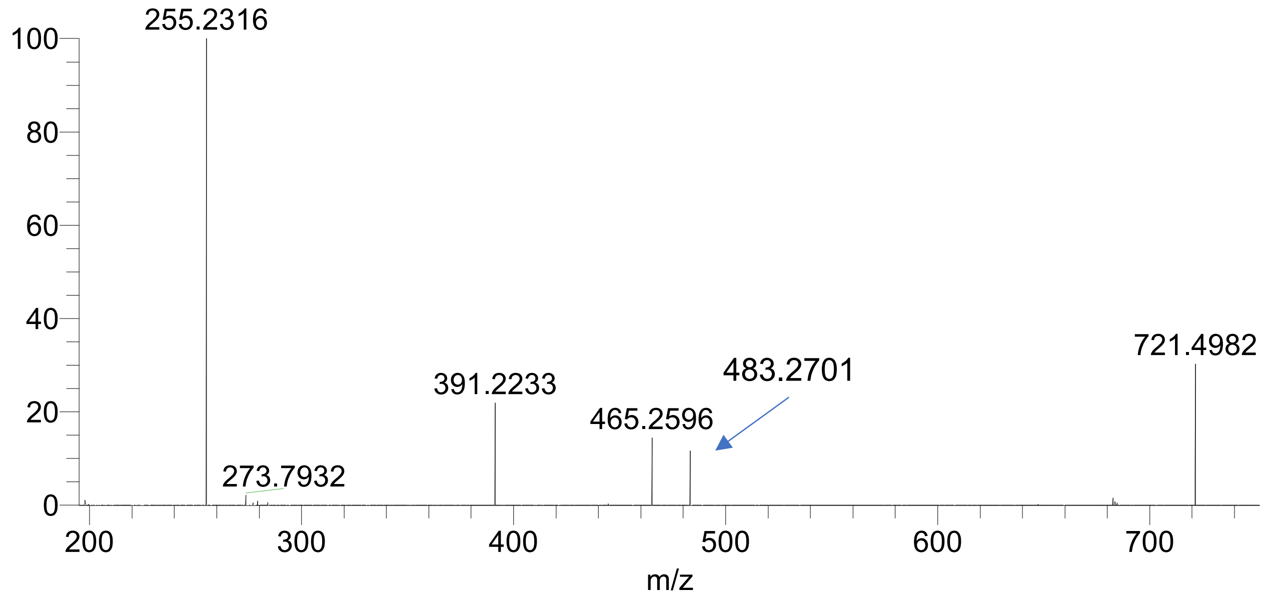


**Supplementary Fig. 30 |** MS/MS spectrum of PG(16:0/16:0) (*m/z* 149.0114) with NCE 28%.

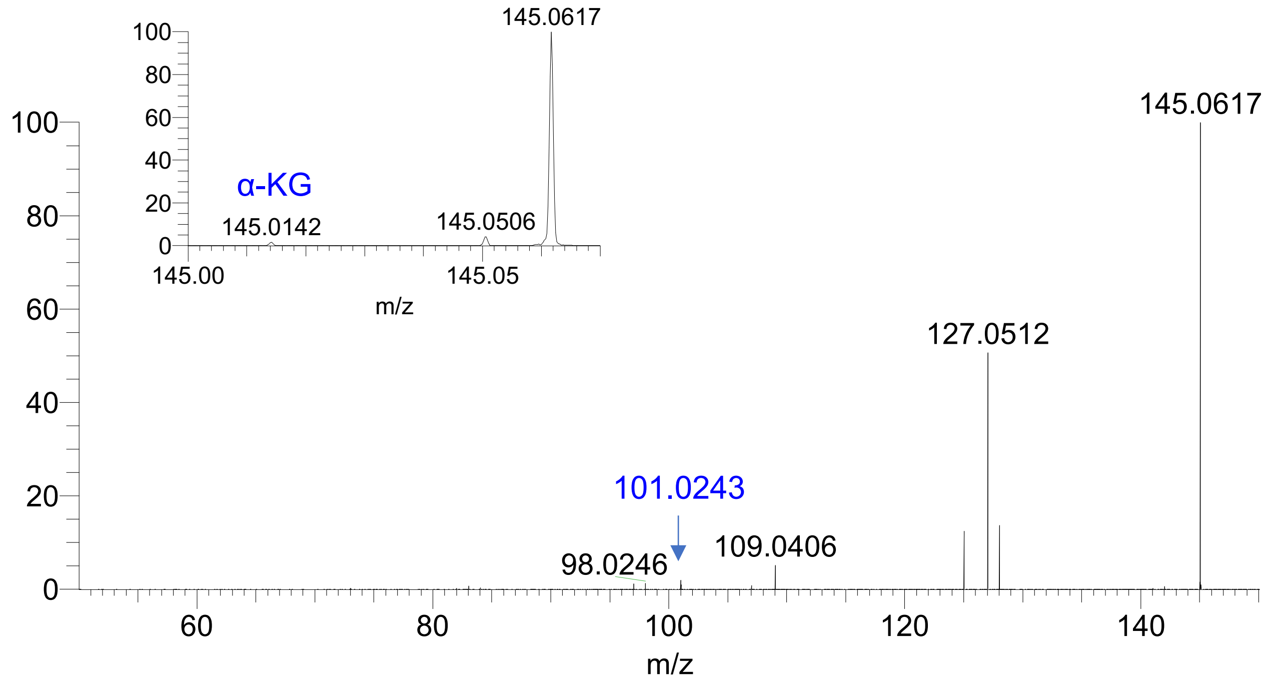


**Supplementary Fig. 31 |** MS/MS spectrum of glutamine (*m/z* 145.0617) with NCE 20%. In addition to glutamine, a small peak from α-ketoglutarate also co-isolated (*m/z* 145.0142) and produced the fragment at 101.0243.

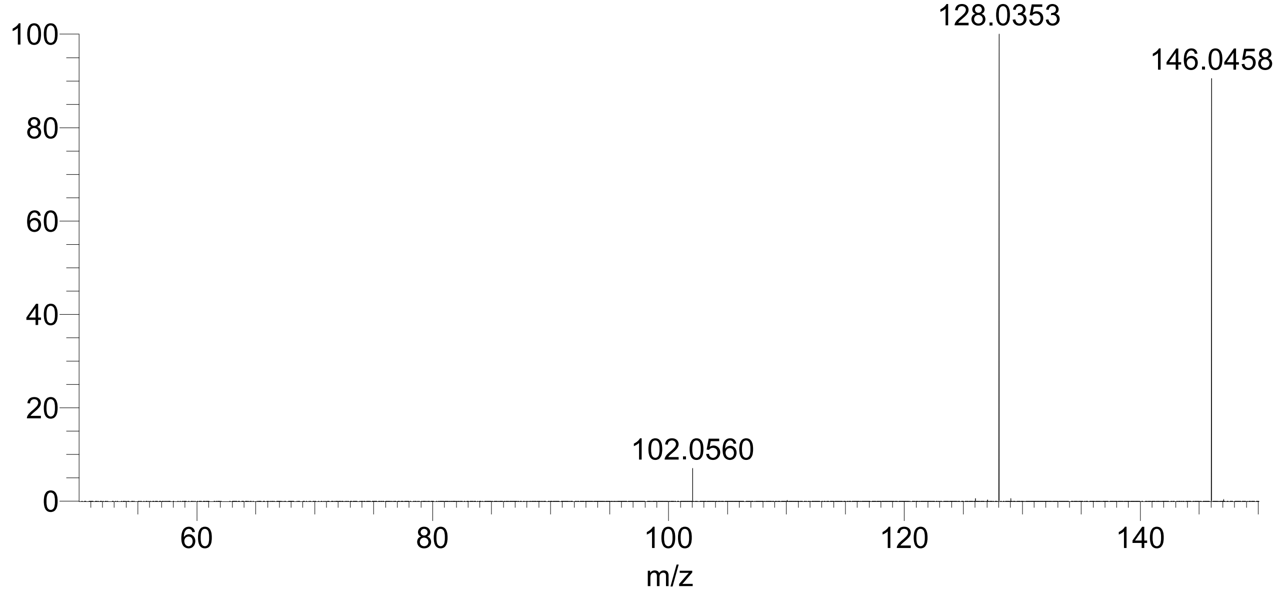


**Supplementary Fig. 32 |** MS/MS spectrum of glutamate (*m/z* 146.0458) with NCE 20%.

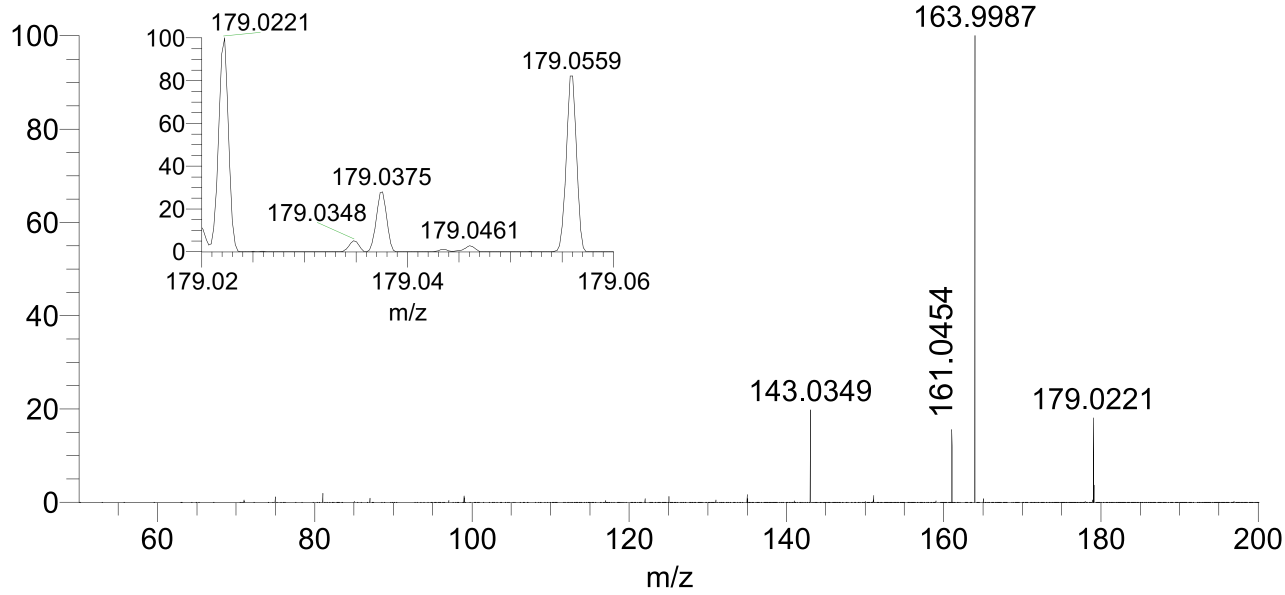


**Supplementary Fig. 33 |** MS/MS spectrum of a hexose (*m/z* 179.0559) with NCE 22%. Fragments *m/z* 161.0454 and 143.0349 are consistent with predictions. The isolation window also captured *m/z* 179.0221, a fragment of HMBOA, as shown by inset.

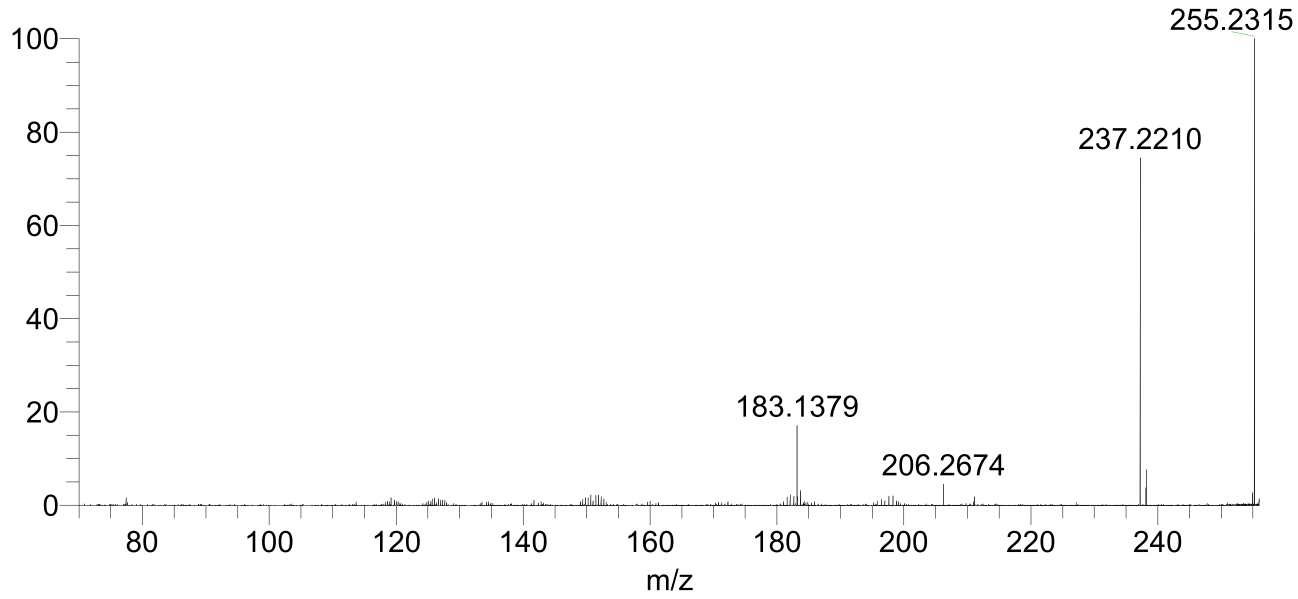


**Supplementary Fig. 34 |** MS/MS spectrum of FA(16:0) (*m/z* 255.2315) with NCE 35%.

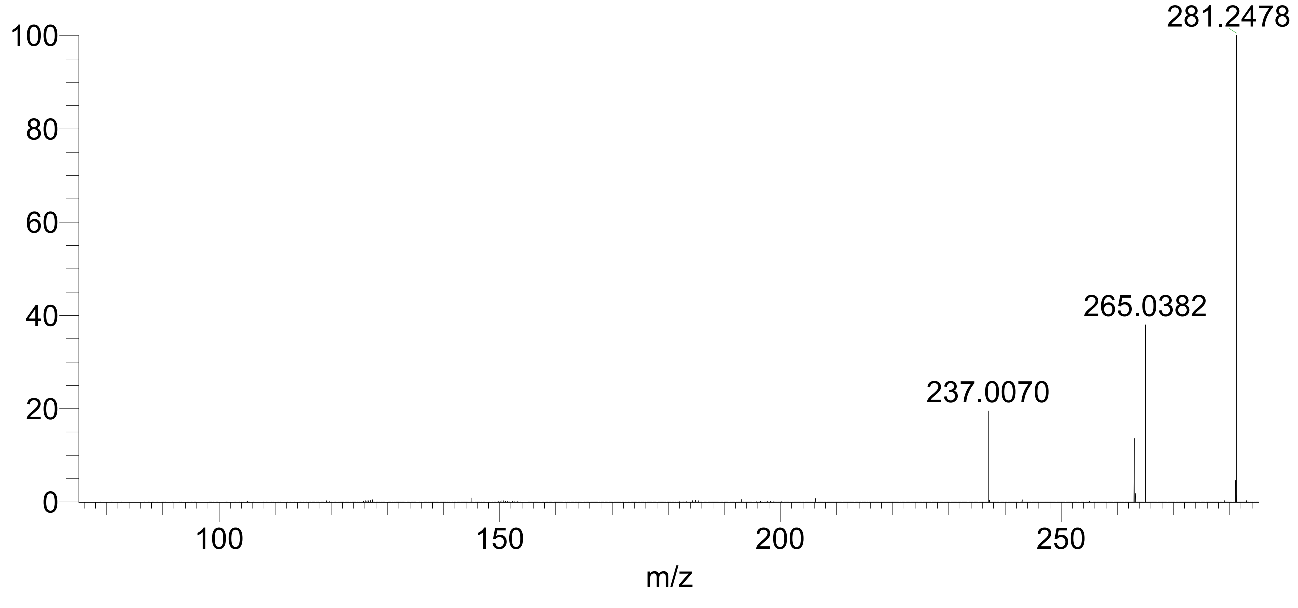


**Supplementary Fig. 35 |** MS/MS spectrum of a FA(18:1) (*m/z* 281.2478) with NCE 35%.
